# Model sharing in the human medial temporal lobe

**DOI:** 10.1101/2021.06.23.449588

**Authors:** Leonie Glitz, Keno Juechems, Christopher Summerfield, Neil Garrett

## Abstract

Effective planning involves knowing where different actions will take us. However natural environments are rich and complex, leading to an exponential increase in memory demand as a plan grows in depth. One potential solution to this problem is to share the neural state transition functions used for planning between similar contexts. Here, we asked human participants to perform a sequential decision making task designed so that knowledge could be shared between some contexts but not others. Computational modelling showed that participants shared a model of state transitions between contexts where appropriate. fMRI data identified the medial temporal lobe as a locus for learning of state transitions, and within the same region, correlated BOLD patterns were observed in contexts where state transition information were shared. Finally, we show that the transition model is updated more strongly following the receipt of positive compared to negative outcomes, a finding that challenges conventional theories of planning which assume knowledge about our environment is updated independently of outcomes received. Together, these findings propose a computational and neural account of how information relevant for planning can be shared between contexts.

## Introduction

Effective goal-directed behaviour requires an agent to learn an accurate model of the world. Theories of reinforcement learning (RL) conceive of this model as a function *p*(*s*′|*s, a*) that encodes the probability of transitioning to a new state, *s*’, given the current state, *s*, and action, *a*. Explicitly learning a state transition function permits agents to plan over possible futures (Sutton and Barto, 1998). This computational framework has been widely used to model simple laboratory behaviours that involve a limited number of state transitions (Daw et al., 2011, Doll et al., 2015, Gläscher et al., 2010, Wunderlich et al., 2012). However, it has well-known limitations, foremost among which is that computational cost grows exponentially with the number of states.

A particular challenge is that in natural environments, transition probabilities can vary with time and context. For example, when planning a journey to work, the likely costs incurred when travelling by car (traffic jams), rail (scheduling delays) or bike (getting wet) can all change unpredictably from day to day. One way to reduce the cost of planning is to share knowledge of likely state transitions across multiple contexts. Consider someone who makes the same journey on a weekday (context 1) or a weekend (context 2). Some knowledge about the transitions that make up the journey may be shared between contexts: if a train line is closed for repairs, then delays can be expected in both circumstances. However, in other cases, knowledge may be specific to contexts. For example, if most drivers on a given route are commuters, then traffic may be heavy on Fridays but light on Sundays. If the brain can learn to use the same transition function in the former but not the latter case, the cost of planning is reduced. Computing invariances over structured state spaces (or cognitive maps) might permit forms of abstract concept formation and inductive reasoning that are thought to support intelligent human behaviour (Behrens et al., 2018).

Here, we developed an experimental paradigm that allowed us to specifically test for “model sharing” – collapsing a state transition function over different contexts. The task required participants to learn a simple associative map in 4 different contexts distinguished via context specific visual cues. In two of the contexts, the same transitions could be used to guide decisions, therefore participants could rely on a shared model, invariant to contextual cues that were present or absent. In the remaining two contexts, model sharing would be detrimental to performance as different state transitions operated independently in each, therefore participants could use the visual cues to distinguish contexts and maintain separate context specific transitions for each.

Our first question was whether, participants would be selective in how they used the contextual cues to filter how information was recruited and updated within and between each context. Our question and approach are similar to those described in a recent paper by Baram and colleagues (Baram et al., 2021), with a key difference being that our work examines how the transition function (how states of the world are associated), rather than the value function (the value of states and actions), is shared between contexts. Our second question was posed at the neural level and addressed by recording fMRI data whilst participants performed the task. We deliberately chose to explicitly instruct our fMRI participants when contexts were coupled and when they were not. This enabled us to detect changes in brain responses at the level of representation that underpinned the large changes in model sharing we observed in participants choices between conditions. Our hypothesis is that similar representational changes would underpin model sharing when this occurs in uninstructed cases (via trial-and-error learning) but caveat that this remains to be tested.

The neural and computational mechanisms by which humans and potentially other animals might share relational knowledge across contexts have come into focus recently, not least because modelling this process is likely to be important for building advanced autonomous agents (Botvinick et al., 2020). A great deal of interest has focussed on the medial temporal lobe (MTL) and adjacent neural structures, which may be important for forming new associations between states (Eichenbaum et al., 1999). For example, repeated co-occurrence of two visual items alters representational properties so that single neurons or neuronal ensembles in the hippocampus and anterior ventral stream come to code for both stimuli (Miyashita, 1988, Yokose et al., 2017, Rey et al., 2018). Moreover, the MTL may be a strong candidate for not only learning state associations in a single context but also encoding relational knowledge that can be generalised across contexts. For example, the MTL is involved in bridging past memories to make new inferences on the basis of paired associations or transitive relations (Bunsey and Eichenbaum, 1996, Wimmer and Shohamy, 2012, Zeithamova et al., 2012, Kumaran et al., 2016, Koster et al., 2018, Park et al., 2019). The MTL is thus an excellent candidate to act as a neural substrate for model sharing, and so we focussed our analyses there.

Finally, we designed our paradigm such that the transition function (the probability of moving from *s* to *s*′ under action *a*) and the value function (when in state *s*, the expected value of taking the action *a* that led to *s*′) were theoretically independent. This was achieved by ensuring that one action (choice of door in our task) led to a high-stakes state where outcomes - signalled in advance - could be either good or bad. An alternate action’s outcomes were always neutral. This allowed us to ask whether state transition learning depends on whether an outcome is positive or negative. From a normative perspective, there is merit in learning about state transitions irrespective of whether they allow you to obtain future benefits or avoid future losses. Classic theories of model based planning thus assume an independence between value and transition functions (Sutton and Barto, 1998). In contrast to this, rodents show greater reverse replay following positive (versus negative) outcomes (Ambrose et al., 2016, Singer and Frank, 2009), an effect that may be the result of positive prediction errors governing the degree to which action policies are worth the effort in updating (Mattar and Daw, 2018). Our paradigm is ideally suited to test whether model updating in humans is symmetric after good and bad experiences.

We tackled these questions using computational modelling and functional neuroimaging. We find that a cluster of regions in the MTL, including the hippocampus, amygdala and entorhinal cortex, display patterns of BOLD activity encoding transition probabilities that are more similar when model sharing between conditions is possible than when it is not. This suggest that the MTL maintains separable encoding patterns for each context in cases where state associations are context specific, but shares encoding patterns when they are not. Next, we show that belief updating of state transition knowledge occurs to a greater degree following positive outcomes compared to negative. This learning asymmetry is reflected by an interaction in the MTL whereby state prediction errors are expressed with greater fidelity for positive compared to negative outcomes. These findings nuance conventional models of planning that assume state transitions and outcomes are tracked and maintained separately from one another.

## Results

### Task and design

Participants (n = 29) performed a planning task in an fMRI scanner (the “heist” task; **Figure 1a**). The task was introduced to participants via a cover story that suggested they were a burglar involved in a heist at one of 4 locations, each denoted by a unique coloured gem. Each trial occurred in one of these four (gem) contexts, and the relevant coloured gem icon remained on the screen throughout the trial to make this clear. After trial onset, participants chose one of two doors (light vs dark), which were respectively associated in context *c* with probabilities *p*_*c*_ of transitioning to the (high-stakes) “heist” state and 1 − *p*_*c*_ of transitioning to the “neutral” state (**Figure 1b**). *p*_*c*_ switched randomly between 0.2 and 0.8 across the course of the experiment, meaning that a door was always likely to transition to one of the outcome states and unlikely to transition to the other. Before making their choice, participants were presented with an additional cue which signalled whether, in the heist state, the participant would be caught (signalled by police cue; incurring a loss) or commit a successful burglary (signalled by swag cue; incurring a gain), whereas no positive or negative outcomes occurred in the neutral state (outcome of zero). The optimal policy was thus to learn the transition probability in order to approach the heist state in the presence of the swag cue and avoid the heist state in the presence of the police cue. To decorrelate choices and probabilities for the scanner, 75% of trials were “forced” in which only a single door was available, but in which transition probabilities could still be updated on receipt of reward. In the remaining 25% of trials, participants could freely choose between the two doors. Participants were unaware during the initial door presentation whether the trial would be a forced or free choice and therefore needed to actively consider transition probabilities on every trial.

**Figure 1.**
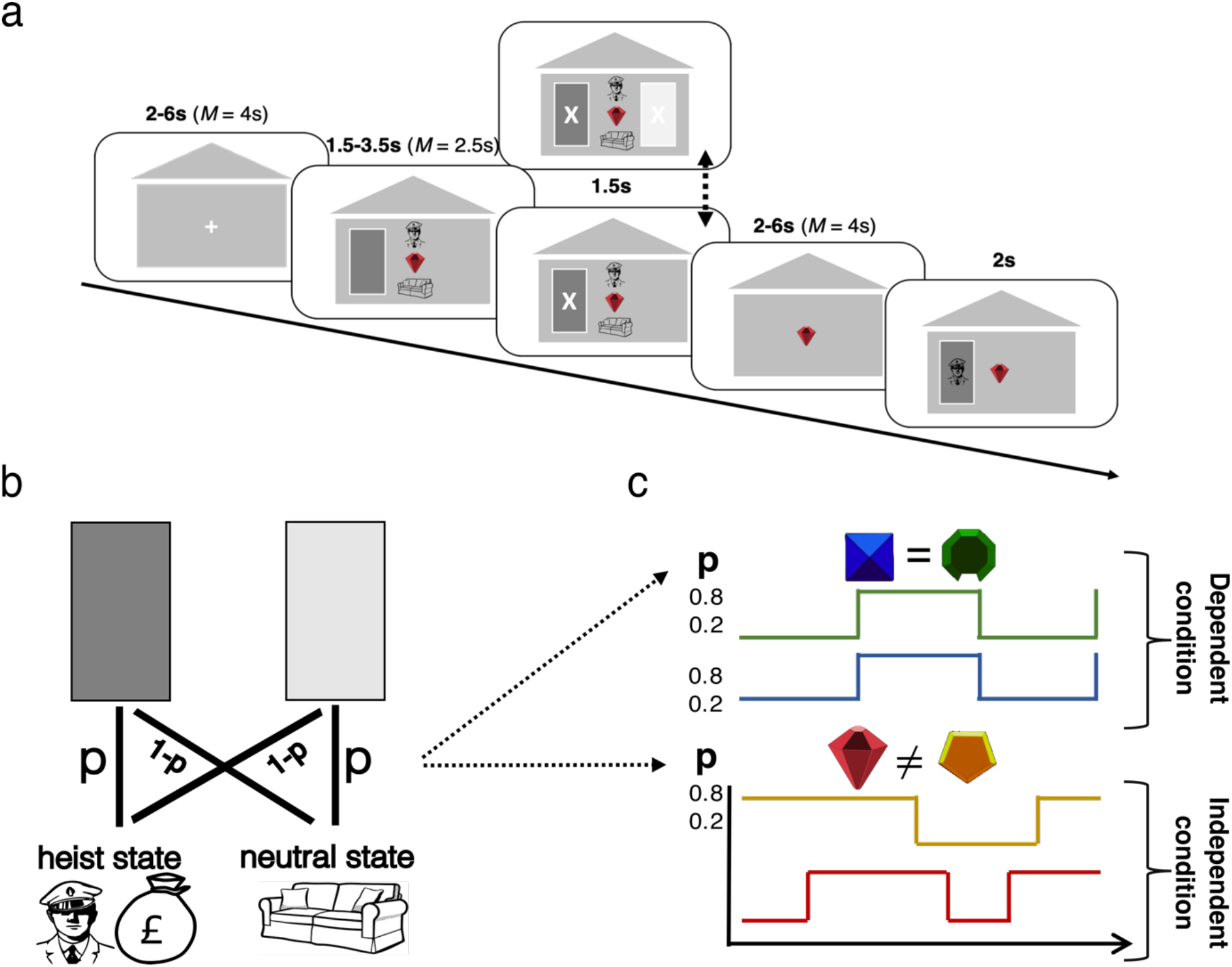
Task Design. **(a)** Trial sequence in the fMRI experiment. On each trial, participants are shown one of two options (a dark and a light door) and either required to select this option (forced choice trials) or to choose between this option and the alternate option (free choice trials). The subsequent state and outcome are then revealed. **(b)** State transition dynamics: at the first stage, each option (framed as two doors) transitions participants to one of two 2^nd^ level states; a “neutral state” in which an outcome of 0 (sofa/chair stimuli) is always obtained, or a “heist state” in which an outcome of 1 (swag bag stimuli) or −1 (police stimuli) can be obtained. One first stage option (light door in the figure) transitions with probability p to the neutral state and with probability 1 - *p* to the vault state; the alternate option (dark door) has the opposite transition probabilities. *p* changes at random points in the task **(c)** In dependent blocks (top panel), *p* is the same in each context; changes to *p* occur simultaneously over the two contexts. In independent blocks (bottom panel), *p* alternates independently in each context. *p* was set to be either .2 or .8 at any given time.

The task was performed in alternating blocks that we label “dependent” and “independent” conditions. n dependent blocks, the transitions probabilities associated with the two contexts (e.g. *p*_1_ and *p*_2_) were yoked so that *p*_1_ = *p*_2_ at all times (**Figure 1c**, top panel). In independent blocks, the transition probabilities associated with the other two contexts (e.g. *p*_3_ and *p*_4_) were unrelated (overlapping on average half of the time; **Figure 1c**, bottom panel). The two contexts that made up each condition were randomly interleaved within a block, but the dependent and independent conditions themselves occurred in temporally distinct blocks of trials. Participants were told before starting the task about the two conditions and were told at the start of each new block whether they were entering a dependent or independent condition block (see **Methods** for full details about the task).

### Behavioural analysis

We first asked whether behaviour differed between the dependent and independent conditions. If participants generalised knowledge of the transition structure across contexts, then they should be more prone to use learning from context *j* to inform subsequent decisions in context *i* when in the dependent than independent condition (note that this behaviour is expected because participants were instructed about the dependence or independence among transition probabilities for the two gems in each block).

We used a logistic mixed effects regression to measure this effect in a trial-history dependent fashion, asking how choices made on each trial *t* in context *i* depended on the history of state transitions observed over the previous 5 trials that had occurred in the contexts *i* and *j*, where *j* was the alternate context within the relevant condition (dependent or independent). To conduct this analysis we recoded choices in a single frame of reference that removed the choice inversion between trials where police and swag cues were present. This was necessary because in our task, the transition history is relevant not for determining the specific response (light vs. dark door) but rather the choice contingent on the presence of the swag or police cue. We call the historic information that is predictive of this recoded choice “transition evidence”.

The results are shown in **Figure 2**. In the dependent condition, transition evidence from the previous two trials (t-2) significantly predicted choice, both when it was experienced in the same (t-1: Fixed Effect β [95% CI] = 1.11 [0.76-1.46], Standard Error (SE) = 0.18, p < 0.001, t-2: β = 0.43 [0.17-0.70], SE = 0.13, p < 0.001) and when it was experienced in the alternate context to the current trial (t-1: β = 0.72 [0.42-1.02], SE = 0.15, p < 0.001, t-2: β = 0.40 [0.17-0.63], SE = 0.12, p < 0.001; **Figure 2a**). In contrast, in the independent condition, choices were only influenced by transition evidence when this was accrued in the same context (**Figure 2b**). This was the case going 1, 2, 3 and 4 trials back (t-1: β = 0.62 [0.24-1.00], SE = 0.19, p = 0.001; t-2: β = 0.30 [0.04-0.55], SE = 0.13, p = 0.02; t-3: β = 0.35 [0.09-0.61], SE = 0.13, p=0.008; t-4: β = 0.32 [0.07-0.58], SE = 0.13, p = 0.01). When transition evidence was accrued in the alternate context, this did not influence participants subsequent choices, even on the immediately previous (t-1) trial (β = 0.07 [-0.12-0.27], SE = 0.10, p = 0.47). See **Supplementary Tables 1** and **2** for full details of all regression coefficients and statistics.

**Figure 2.**
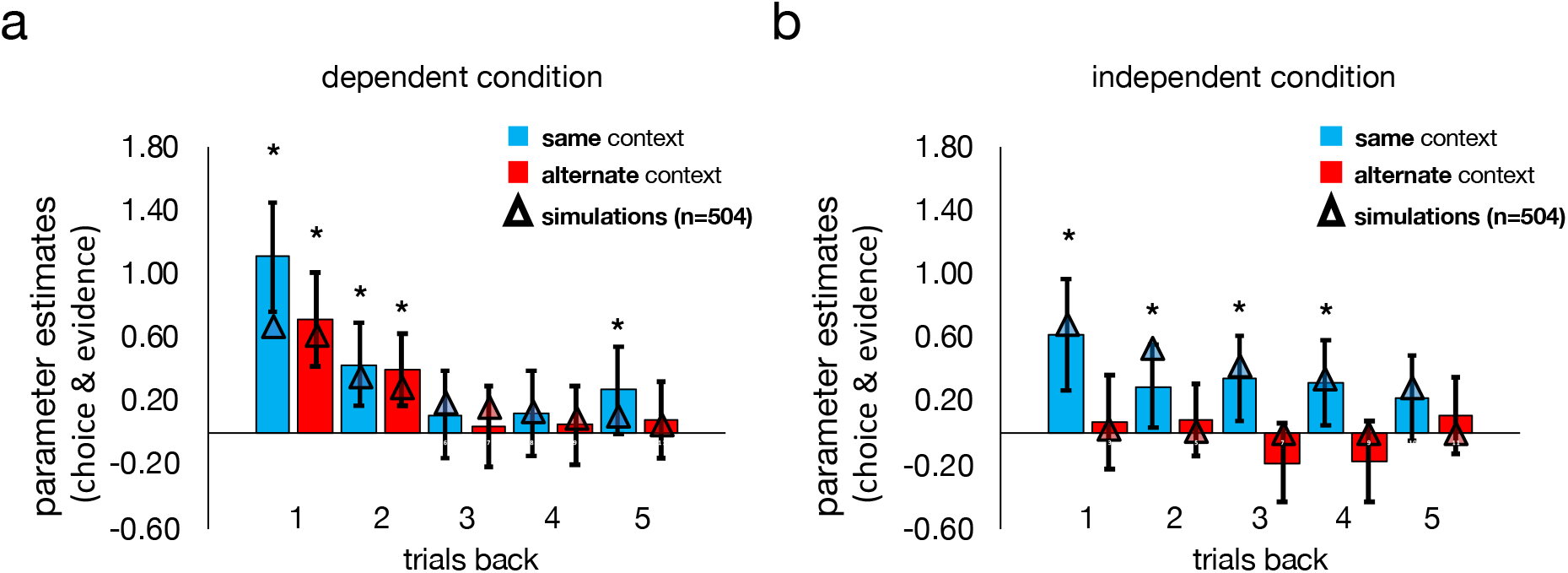
Behavioural data. Parameter estimates predicting choice from state transitions experienced 1-5 trials back, separated according to whether transitions occurred in the same (blue) or alternate (red) context to the current trial context in **(a)** the dependent condition and **(b)** the independent condition. Bars represent fixed effects regression coefficients from a mixed effects logistic regression on participants’ choices. Triangles represent the mean fixed effects regression coefficient estimates generated via the same mixed effects logistic regression as the data but for choice simulated for agents under a flexible computational learning model which enables evidence integration to adapt to the condition in which choices are being made (dependent or independent). *p < 0.05 (human data). Error bars express 95% confidence intervals.

To directly compare the relative weight participants placed on past evidence received from the same and alternate context in each of the two conditions (dependent, independent) we ran an additional mixed effects model. We computed the difference in transition evidence between the two contexts (averaged over the past 5 trials; we call this “differential evidence”) for each condition (dependent/ independent) and their interaction as predictors in this model. This revealed a significant interaction between differential evidence and condition (β = −0.41 [-0.63, −0.18], SE = 0.11, p < 0.001) along with a main effect of differential evidence (β = 0.43 [0.20, 0.66], SE = 0.12, p < 0.001), but no main effect of condition (β = −0.05 [-0.13, 0.04], SE = 0.04, p = 0.27). The interaction between differential evidence and condition remained significant in a permutation test which guards against greater similarity of feedback (in the dependent condition compared to the independent condition) confounding the effect (β = −0.41, 95% range under the null distribution: [-0.25, 0.08], p < 0.001). Together, these results suggest that the relative preference for information received from the same (versus the alternate) context shifted between conditions. This was a result of participants increasing integration of information from the alternate context in the dependent condition.

We also analysed data from an additional pilot experiment (n = 31; see **Methods**) with an identical structure except for two important differences: firstly, there were no forced choice trials, and secondly, participants were not instructed about the dependence or independence of the transition structure but were left to discover it for themselves. In contrast to the fMRI cohort, in the independent condition, choices were influenced by transition evidence that accrued in *both* the same context (t-1: β [95% CI] = 1.20 [0.95, 1.45], SE = 0.13, p < 0.001, t-2: β = 0.65 [0.49, 0.82], SE = 0.08, p < 0.001, t-3: β = 0.39 [0.26, 0.52], SE = 0.07, p<0.001, t-4: β = 0.25 [0.15, 0.34], SE = 0.05, p<0.001, t-5: β = 0.19 [0.09-0.29], SE = 0.05, p<0.001) and in the other context (t-1: β = 0.22 [0.07, 0.37], SE = 0.08, p = 0.004, t-2: β= 0.15 [0.06, 0.24], SE = 0.05, p = 0.001, t-3: β = 0.10 [0.002, 0.20], SE=0.05, p=0.046, t-4: β = 0.09 [0.004, 0.179], SE=0.04, p=0.04, t-5: β = 0.09 [-0.001, 0.19], SE=0.049, p=0.05). See **Supplementary Figure 9** and **Supplementary Tables 9** and **10** for plots and full details of regression coefficients and statistics. Examining whether differential evidence interacted with condition revealed a significant interaction between differential evidence and condition (β = −0.097 [-0.17, −0.02], SE = 0.04, p = 0.010) however this was not significant in the permutation test (β [95% range under the null distribution] = −0.097 [-0.25, 0.08], p=0.92). As discussed below, we think that together these results suggest that in the absence of instruction, participants may have a stronger prior that the two gems which co-occur in time belong to a shared latent context.

### Computational model

Our modelling framework assumed that choices were determined by a mixture of associations learned in an independent and dependent fashion across contexts. We note at the outset that our model is not intended primarily as an account of the computations that humans undertake, but as an analytic tool that compactly parameterises human policy with just a few parameters which allows us to verify the degree to which humans share a model between contexts.

The model is composed of two learners, one which learns a shared transition function across a pair of contexts and another which learns a separate transition function for each context. On each trial, choices are determined by linearly mixing the estimated probabilities from each learner according to a weighting parameter *w*, and using the resulting probabilistic estimate 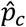 to compute the relative expected value of heist and neutral states, according to which a choice was made via inverse temperature parameter *β*. The optimal policy (for an omniscient agent) would be to use *w* = 1 in the dependent condition and w = 0 in the independent condition. On each trial, participants updated the context specific and the context independent transition functions according to a state prediction error, δ, which quantifies the degree of surprise at reaching a state given the option chosen and current estimates of the transition function. δ was also weighted by *w* and the degree of update governed by a learning parameter, α.

We compared two versions of this learning model. A *fixed model* in which *w* was held constant across the experiment was compared to a *flexible model* in which *w* was allowed to reverse between experimental conditions (i.e., ***w***_independent_ = 1 - ***w***_*dependent*_). This feature of the flexible model gives it the capacity to shift between relying to a greater degree on separate transition functions in the independent condition (i.e., *w* towards 0) and relying on a shared transition function in the dependent condition (i.e., *w* towards 1). See **Methods** for full model specification.

### Flexible model adapts information integration between conditions

We fit each model to single subject choices on a per-trial basis and compared fixed and flexible models by computing unbiased marginal likelihoods via subject-level Leave One Out cross validation (LOOcv) for each participant. Comparison of LOOcv scores revealed significantly lower scores (indicating superior performance in cross validation at predicting participant choices) for the flexible model compared to the fixed model (t(28) = 2.72, p < 0.01, see **Table 1** for model parameters and LOOcv scores). The best-fitting *w* parameter tended towards 1 in the dependent condition and 0 in the independent condition, consistent with the behavioural data. This indicates that participants learned a single transition function in dependent blocks but reverted to learning two different transition functions in independent blocks (by contrast, when *w* was held fixed across blocks it assumed an intermediate value of ∼0.61). A flexible model with two separate *w* parameters (one per condition, fitted separately) did not account any better for participants choices than the flexible model with a single *w* that reversed between conditions (t(28) = −1.59, p = 0.12). Simulating choices using a population of subjects drawn according to best-fitting parameters of the flexible model showed that the flexible model qualitatively recapitulated the change in relative preference for information from the alternate versus same context between conditions (**Figure 2**). The fixed model was not able to recover this pattern (**Supplementary Figure 1**).

**Table 1.** Model fitting and parameters. The table summarizes for each model its fitting performances and its average parameters: LOOcv: leave one out cross validation scores, mean (standard error of the mean) over participants; free parameters: number of free parameters in each model; *α*: learning rate; *β*: softmax slope (sensitivity to the difference in the value of choosing dark versus light door (on free choice trials). *w*: weighting parameter (governs the weighted combination of context independent and context dependent transition functions). Data for model parameters are expressed as mean and 95% confidence intervals (calculated as the sample mean +/- 1.96 x standard error). *p < 0.01 comparing LOOCV scores between the two models (paired sample t-test). Lower scores indicate superior performance in cross validation.

| | LOOcv scores | free parameters | $\beta$ | $\alpha$ | $w$ (fixed) | $w$ (dependent condition) | $w$ (independent condition) |
| --- | --- | --- | --- | --- | --- | --- | --- |
| fixed model | 47.47<br>(+/- 1.26) | 3 | 1.80<br>[95% CI = 1.40, 2.20] | 0.65<br>[95% CI = 0.47, 0.80] | 0.61<br>[95% CI = 0.49, 0.70] |  |  |
| flexible model | 46.32*<br>(+/- 1.47) | 3 | 1.59<br>[95% CI = 1.24, 1.95] | 0.56<br>[95% CI = 0.38, 0.73] |  | 0.85<br>[95% CI = 0.66, 0.95] | 0.15<br>[95% CI = 0.05, 0.34] |

### Neuroimaging data

Having established that participants behave differently in the dependent and independent conditions, we turned to the fMRI data to understand the neural mechanisms that supported this differential behaviour. Our goal was to use multivariate approaches (including representational similarity analysis, RSA) to examine how multivoxel patterns encoding transition probabilities (i.e. beliefs about the forthcoming state) were related in the dependent and independent conditions. However, we first adopted a univariate analysis to identify target sites for the coding of the state transition function, using the state prediction error (SPE) from the model. We expected that the MTL would be sensitive to SPEs, consistent with a long tradition implicating the hippocampus in the formation of state associations (Eichenbaum et al., 1999), and a detector of states that either match or violate the agent’s expectations (Kumaran and Maguire, 2007, Duncan et al., 2012).

### Univariate analysis

We thus modelled BOLD responses at the time the transitioned-to state (“heist” or “neutral”) was revealed using a parametric predictor encoding the unsigned state prediction error |δ| extracted from the flexible model. This analysis collapsed over conditions (dependent, independent). This modulator was included alongside other quantities coding for outcome, trial type (forced/free choice) and the interaction of outcome and |δ|(see **Methods**).

The BOLD signal correlated negatively with |δ| in two MTL clusters (peak left: [x, y, z]: −20, −4, −28, t(28) = 5.01, p < 0.001 uncorrected for multiple comparisons; peak right: 18, −4, −21, t = 4.22, p < 0.001 uncorrected), which survived small volume correction using a bilateral anatomical MTL mask (peak left: −20, −7, −24; t(28) = 5.01, family wise error corrected at the peak level within bilateral MTL mask [p_FWE_] = 0.008; peak right: 18, −4, −21; t(28) = 4.22, p_FWE_ = .047). In total, 10.14% of voxels lay within the anatomically defined amygdala, 33.33% within the hippocampus, 49.28% within the parahippocampus and 5.80% in the entorhinal cortex (**Figure 3a**), determined by assessing overlap with anatomical masks generated in WFU pickatlas (see **Methods**).

**Figure 3.**
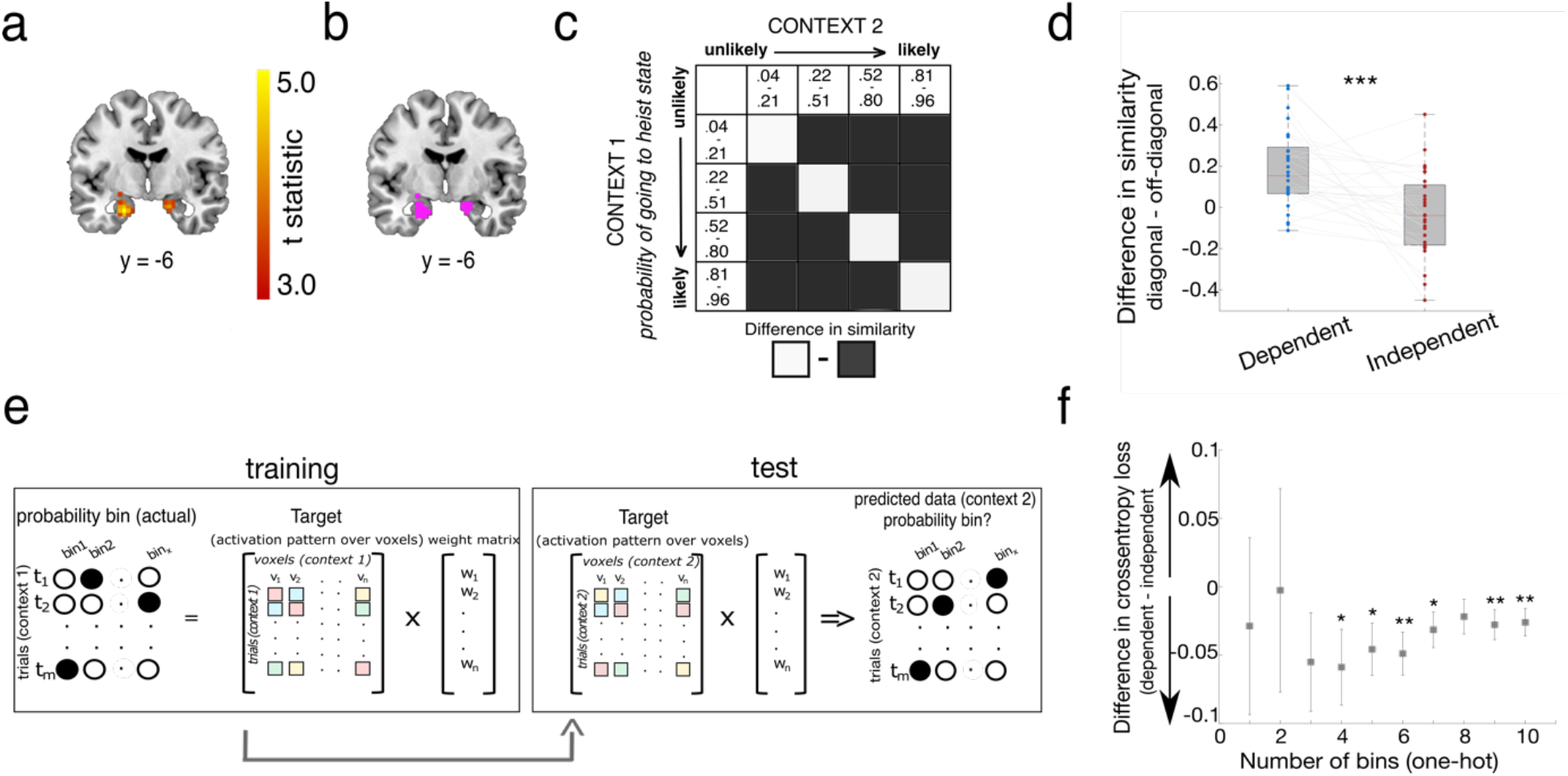
**(a)** The magnitude of (unsigned) state predictions errors related negatively to the degree of BOLD response in bilateral MTL. Image shown thresholded at p < 0.001 uncorrected. **(b)** Voxels in this contrast were converted to a bilateral mask and used as a functional ROI in subsequent analysis. **(c)** Schematic of the RSA analysis at the time of planning (door presentation). In each context, trials were divided into quartiles, according to participants’ current estimates of p (heist state | door presented) extracted from the computational learning model (mean quartile ranges: bin 1: *p* <= 0.21, bin 2: 0.21 < *p* <= 0.51, bin 3: 0.51 < *p* <= 0.80; bin 4: 0.80 < *p* <= 0.96). **(d)** Difference scores were significantly greater for dependent than independent blocks. Dots represent individual participant data, grey lines indicate datapoints belonging to the same participant. Red line indicates the median, box represents the 25^th^ and 75^th^ percentile of data, whiskers extend to any data point that is not outside 1.5 times the interquartile range **(e)** Schematic of the encoding model analysis (example shown for one-hot case). **(f)** Difference in cross-entropy loss from the encoding model between dependent and independent blocks (predicting probability bins in one context in a condition using weights trained on the other context in that condition; in crossvalidation) for a range of probability bins (one hot case). Error bars show standard error of the mean; *\* significant at p < 0*.*05, ** significant at p < 0*.*01*., *\*\*\* significant at p < 0*.*001*

The negative direction of the parametric effect indicates greater BOLD in response to expected (compared to unexpected) state transitions as previously observed in other learning tasks at the time new information is received and beliefs are updated (Sharot et al., 2011). We combined these clusters (extracted at p < 0.001 uncorrected) into a single bilateral functional region of interest (ROI) mask (**Figure 3b**) which we then used for subsequent multivariate analyses.

### Representational Similarity Analysis (RSA)

Next, we used a multivariate approach to assess the mapping from BOLD responses in our functional ROI to transition probabilities, and to measure how this mapping changed over contexts. We began with an analysis of BOLD signals at the time of choice, i.e. when the door was presented. This is the timepoint during which participants needed to consider the transition probability to each prospective 2^nd^ level state. We first used representational similarity analysis (RSA), measuring the correlation distance across multivoxel patterns associated with transition probabilities *p*(heist state | door presented) derived from our flexible learning model into quartiles, both across blocks and across gems (**Figure 3c**). Note that our prediction is that neural patterns encoding transition probabilities should be more similar across contexts in dependent than in independent blocks. We thus computed a similarity score by averaging correlations in diagonal (same probability quartile) vs. off-diagonal (different probability quartile) cases, separately for the two contexts in the dependent and independent condition (**Figure 3**).

This revealed a significant condition (dependent, independent) x quartile (diagonal, off-diagonal) interaction (t(28) = 4.02, p < 0.001, 95% CI [.11,.33], paired sample t-test). This was the result of a difference in similarity between on and off diagonal scores in the dependent condition (t(28) = 5.33, p < 0.001, 95% CI [.11,.26], one sample t-test versus 0) which was absent in the independent condition (t(28) = −0.82, p = 0.42, 95% CI [-.11, .05], one sample t-test versus 0, see **Figure 3d**).

One interpretation of this finding is that in the dependent condition, the MTL encodes the state transition function for each context with a common neural pattern. However, we also considered some alternative possibilities. First, we examined whether the results held if we allocated trials to bins using fixed probabilities across the unity range (i.e., quartile 1: 0.00-0.25, quartile 2: 0.26-0.50, quartile 3: 0.51-0.75 & quartile 4: 0.76-1.00) rather than adapting bins for each participant according to the specific distribution of probabilities they used. This revealed the same pattern of results (condition*diagonal interaction: t(28) = 4.20, p < 0.001; 95% CI [.10, .30]; difference in similarity between on and off diagonal bins in the dependent condition: t(28) = 5.73; p < 0.001; 95% CI [.13, .28]; difference in similarity between on and off diagonal bins in the independent condition: t(28) = 0.04; p = 0.97; 95% CI [-.07, .08], see **Supplementary Figure 4**). Second, we checked that the number of trials in each probability quartile were well matched between contexts, finding that they were (t(28) = 1.50, p = 0.55, 95% CI [-.02, .15], see **Supplementary Table 3**). Finally, we were concerned that the effect might arise as a spurious effect of closer temporal proximity between trials in the same transition probability quartile in dependent blocks. To address this, first we checked whether the average difference between the temporal distance of trials in on versus off diagonal quartile combinations was correlated with the difference in representational similarity (see **Methods** and **Supplementary Figure 6**). This was neither the case in the dependent condition (r = -.20, p = 0.32), nor the independent condition (r = .15, p = 0.44). As an additional check, we then repeated our analysis in cross-validation across sessions. In other words, we measured the similarity between quartile/bin *n*_*i*_ and *n*_*j*_ where *i* and *j* are drawn from different scanner runs and computed the average for each similarity bin across all possible *c*_1*i*_ and *c*_2*j*_ combinations where *i* ≠ *j* (see **Supplementary Figure 8** for illustration). This revealed the same (albeit weaker) pattern of results with fixed probability bins (condition*diagonal interaction: t(28) = 2.04, p = 0.05; 95% CI [-.00, .06]) and probability quartiles (t(28) = 1.89, p = 0.069; 95% CI [-.00, .07]).

### Multivariate encoding model

Next, taking a complementary approach, we built an encoding model that mapped transition probabilities (in the frame of reference *p(state = heist*|*door presented)* derived from the flexible learning model as before) flexibly onto voxels within the MTL ROI, separately for each context *c*_*b*_. We then inverted this model to predict transition probabilities both for the same context *c*_*b*_ and the other three (held out) contexts (contexts) *c*_*b*_ where *a* ≠ *b* (see **Figure 3e** for a schematic of this analysis). This approach allowed us to train and test in cross-validation, by obtaining weights from session (scanner run) *i* and then using these to predict the probabilities for each context on session *j*. The model output was a 4 × 4 (context × context) matrix of predicted vs. true (model-derived) transition probabilities, which we compared via cross-entropy loss. This allowed us to measure whether, within the MTL, neural patterns coding for probabilities were more similar across contexts in the dependent condition (e.g. *c*_1_ → *c*_2_ and *c*_2_ → *c*_1_) than in the independent condition (e.g. *c*_3_ → *c*_4_ and *c*_4_ → *c*_3_). Unlike the RSA approach, this also allowed us to compare two different coding schemes. It could either be the case that state associations are encoded in a high-dimensional format in which probabilities map onto bins with no input structure. This can be implemented via a one-hot input function in the encoding model which also enables us to test various levels of granularity of binning, to verify that the RSA results were not specific to our choice of having 4 bins. Alternatively, it could instead be the case that probabilities are encoded in a low-dimensional format, whereby neural patterns are more similar for closer probabilities (e.g., bin 1 is more similar to bin 2 than to bin 4). This can be implemented via an Gaussian input function (effectively, a tuning curve for probability) in the encoding model. Probabilities were converted to odds ratios for this exercise (see **Methods**).

The results validated and extended those of the RSA. Using one-hot encoding of probability, we found stronger evidence of shared encoding of probability in the dependent condition compared to the independent condition. Furthermore, this effect was independent of the number of bins chosen, as long as there were more than 3 bins (**Figure 3f**). We obtained the most robust effects with ∼6 bins (implying a psychologically plausible granularity to the estimation of transition probabilities), for which the cross-validated loss was substantially higher between contexts in the independent condition than those in the dependent condition (t(28) = −3.12, p = 0.002). When cross-validation was performed across sessions only, reconstructing both probabilities in the other context as well as in the same context using information from another session (e.g. *c*_1_ *session* 1 → *c*_1_ *session* 2) we found the same pattern of results (t(28) = −3.80, p < 0.001. Similar results were also obtained at different granularities. Interestingly, we were unable to recreate these effects under the additional constraint imposed by Gaussian encoding of probability ratios. This implies that whilst there is a consistent code for transition probabilities, its similarity structure does not map smoothly onto the one-dimensional axis given by probability.

### Characterising the nature of the effect in the medial temporal lobe

Our functional ROI (**Figure 3b**) used for the RSA and encoding model analysis included voxels from different anatomical subregions of the MTL. To investigate whether the observed effect was specific to particular subregions of the MTL, we conducted four further RSA analyses. This was exactly as described above but on voxels from 4 different MTL anatomical masks (see **Methods**): hippocampus, parahippocampus, entorhinal cortex and amygdala (**Figure 4a**). Fisher transformed similarity scores were then entered into a region by condition (dependent/ independent) 4*2 repeated measures analysis of variance (ANOVA).

**Figure 4.**
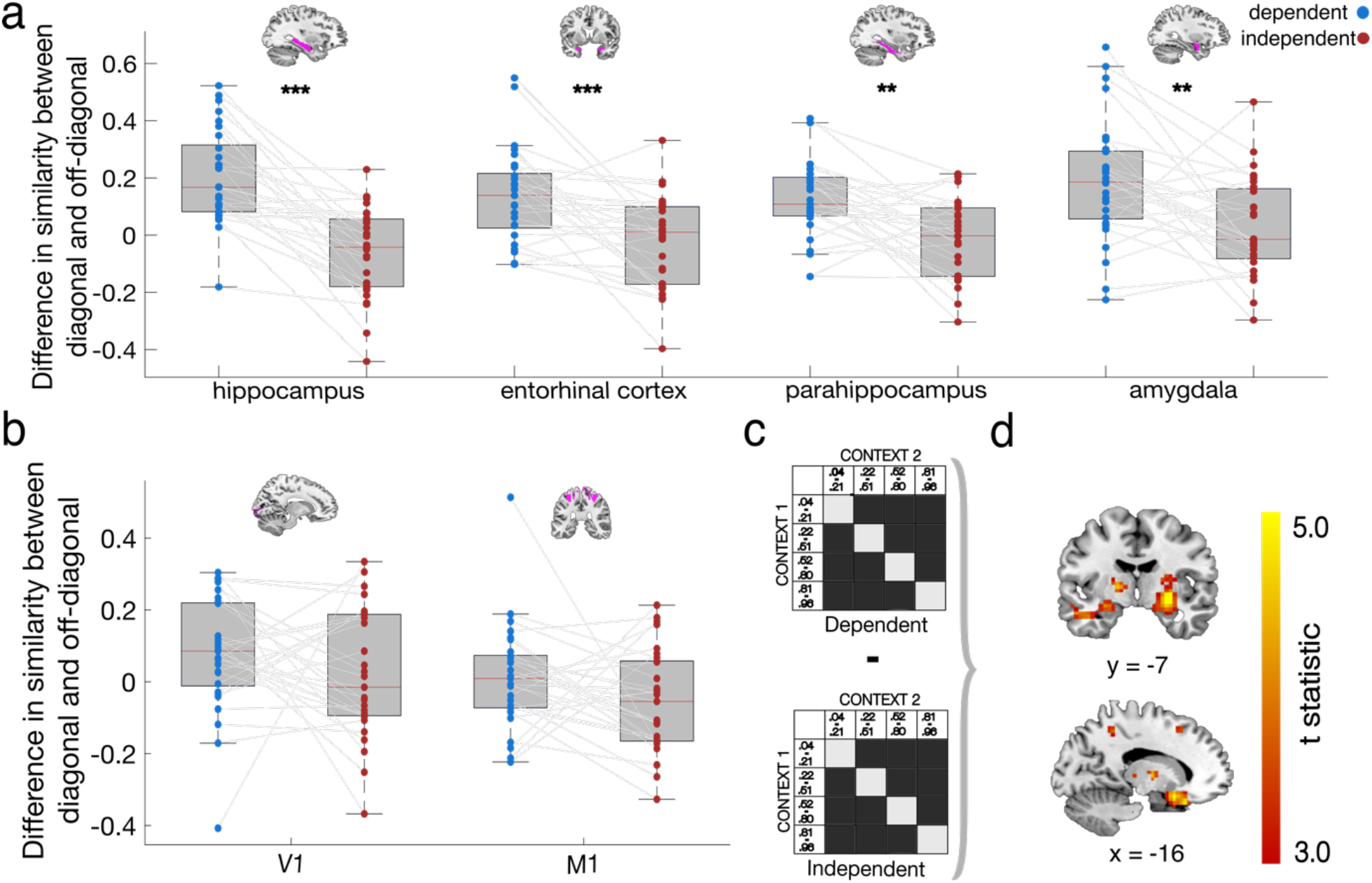
RSA analysis in **Figure 3c** was repeated using **(a)** anatomical masks of subregions of the MTL, specifically: bilateral hippocampus, parahippocampus, entorhinal cortex and amygdala, **(b)** as well as in V1 and M1 (control regions). **(c)** illustration of the whole brain searchlight interaction analysis; the difference between on diagonal and off-diagonal similarity was contrasted between conditions. **(d)** whole-brain searchlight interaction analysis revealed greater similarity between on versus off diagonal in the dependent condition compared to the independent condition in our functional ROI, right dorsal striatum (top panel) and left inferior frontal gyrus/OFC (bottom panel). Brain images shown at p < 0.001 uncorrected, thresholded at t > 3.

This revealed a main effect of condition (F(1,28) = 29.40, p < 0.001) with the difference in similarity (M) between on and off diagonal scores greater in the dependent than independent condition (M_hippocampus_ = .26, M_parahippocampus_ =.13, M_entorhinal cortex_ = .15, M_amygdala_ = .16) as well as a region × condition interaction (F(2.45, 68.54) = 3.91, p = 0.018; Greenhouse-Geisser corrected). To better understand the interaction we proceeded to test the difference in similarity scores between conditions in each region with every other region (correcting for multiple comparisons). This revealed a larger difference between conditions in the hippocampus compared to each of the other 3 MTL subregions (entorhinal cortex, amygdala and parahippocampus, all p < 0.05, paired sample t-test) with the parahippocampus surviving correction for multiple comparisons (t(28) = 3.91, p = 0.001, significant at Bonferroni-corrected threshold of p < 0.008; for full table of paired comparisons see **Supplementary Table 5**). There was also a main effect of region (F(3, 84) = 3.33, p = 0.023), with the difference across both conditions being significantly greater in the amygdala than in both parahippocampus (t(28) = 3.07, p = 0.005) and entorhinal cortex (t(28) = 2.81, p = 0.009). Together, these results suggest that greater similarity in transition encoding in the dependent compared to the independent condition was not exclusive to a particular subregion of the MTL but was most pronounced in the hippocampus (see **Figure 4a** and **Supplementary Table 5**).

Finally, to test if the differences between our conditions was selective to the MTL or present over the whole brain, we conducted the same RSA analysis using voxels in two control regions: early visual cortex (V1) and primary motor cortex (M1). There was no significant difference between conditions in either control region (V1: t(28) = −0.28, p = 0.78, 95% CI [-.08,.06]; M1: t(28) = −0.22, p = 0.83, 95% CI [-.09,.08], **Figure 4b**).

### RSA whole-brain searchlight

Next, we repeated the same RSA as described above across the whole brain using a searchlight approach. In the dependent condition, this identified activity within our functional ROI (right peak: 22, −7, −18; t(28) = 4.31, family-wise error corrected at peak level within functional ROI mask, p_FWE_ = 0.005; left peak: −24, −7, −18; t(28) = 4.92, family-wise error corrected at peak level within functional ROI mask, p_FWE_ = 0.001). The cerebellum also survived family wise error correction for multiple comparisons at the cluster level (cluster-defining threshold p < 0.001, uncorrected; see **Supplementary Material**). We did not find any evidence for differences in similarity in or outside our functional ROI in the independent condition, even at very lenient thresholds (p < 0.01 uncorrected).

Next, we conducted a new searchlight that directly tested the difference in similarity for on versus off diagonal bins between conditions (**Figure 4c**). This also revealed activation in our functional ROI (right peak: 22, −7, −18; t(28) = 3.55, family-wise error corrected at peak level within functional ROI mask, P_fwe_ = 0.02; left peak: −27, −10, −14; t(28) = 4.02, family-wise error corrected at peak level within functional ROI mask, p_FWE_ = 0.006). A cluster in the right dorsal striatum (29, −7, −7: t(28) = 5.80, p < 0.001, FWE cluster-level corrected, **Figure 4d**) which extended into the hippocampus, as well as the inferior frontal gyrus adjacent to Brodmann area 47 (−16, 14, −18: t(28) = 5.08, p = 0.007) and left cerebellum (−2, −46, −21: t(28) = 4.92, p = 0.03) also survived family wise error correction for multiple comparisons over the whole brain at the cluster level (cluster-defining threshold p < 0.001 uncorrected).

### Multivariate analysis during transition probability updating

The analyses described so far focus on the timepoint when planning takes place (door presentation). What happens during updating? To examine this, we conducted a related analysis at the time of transition outcome, i.e. when participants learned whether, conditional on their choice, they had reached the “heist” or the “neutral” state. We reasoned that in order to update the state action representations appropriately (in a shared or unshared manner across contexts) it would be necessary to re-encode both the selected action (light vs. dark door) and encountered state (heist vs. neutral). We thus partitioned data according to these factors and investigated whether BOLD signals were more similar when both state and action matched (i.e. on-diagonal elements) vs where they did not (off-diagonal elements) at the time of updating, separately for the dependent and independent conditions (**Figure 5a**) in our functional ROI. This analysis also revealed a significant condition × diagonal interaction (t(28) = 2.67, p = 0.01 95% CI [.03,.22], paired sample t-test, **Figure 5b**), driven by a significant difference in similarity between matched and mismatched choice-state combinations (the on versus off diagonal in **Figure 5a**) in the dependent condition (t(28) = 2.35, p = 0.03 95% CI [.01, .18], one sample t-test versus 0) which was absent in the independent condition (t(28) = - 0.78, p = 0.44, 95% CI [-.12, .05]).

**Figure 5.**
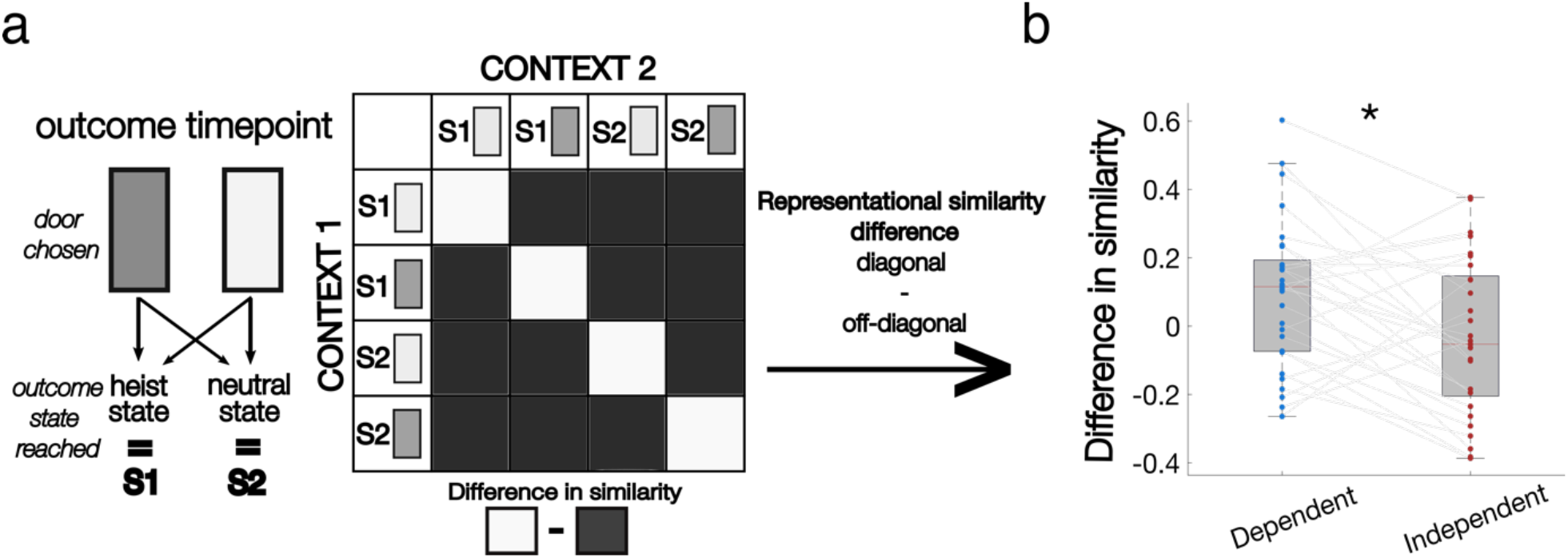
**(a)** Model representational dissimilarity matrix. We examined the BOLD similarity at the time of outcome between matched choice (door) – outcome state combinations and mismatched combinations between contexts in the two conditions. **(b)** In our MTL ROI, the difference between representational similarity of matched and mismatched combinations was significantly greater in dependent than independent blocks.

Once again, this effect was not specific to a particular subregion of the MTL (see **Supplementary Figure 7**). We entered similarity scores from RSAs conducted in subregions of the MTL into a condition (dependent, independent) by region (hippocampus, parahippocampus, entorhinal cortex, amygdala) ANOVA. This revealed a main effect of condition (F(1,27) = 10.05 p = 0.004). There was no main effect of region (F(3,81) = 0.95, p = 0.42), nor a region by condition interaction (F(3,81) = 1.21, p = 0.31; see **Supplementary Figure 7**). There was no difference between conditions in two control brain regions (V1: t(28) = 1.16, p = 0.26, 95% CI [-.04,.16]; M1: t(28) = 1.66, p = 0.11, 95% CI [-.02,.15]). See **Supplementary Figure 7** for whole brain searchlight results.

### Outcome valence modulates updating of state transitions

An interesting feature of our design is that the transition function changes (with reversals of *p*) in a way that is unrelated to outcomes. This means that in theory, any learning about the transition function should not depend on whether the outcome was positive or negative. To test whether participants might be biased to update the transition function more or less according to the outcome, we calculated a ***consistency score*** (see **Methods**) for each participant. This measured the consistency of each choice given transitions experienced on the previous trial. A high consistency score indicates that a participant updates transitions strongly on the basis of feedback. This was calculated separately for trials in which participants received a positive outcome (+1 on a gain trials or 0 on a loss trial) and those where they received a negative outcome (−1 on a loss trial or 0 on a gain trial) on the previous trial. Notably, this is not the same as a win-stay, lose-switch bias, as a choice would be considered consistent only if it considered *both* past transitions and the current reward/loss incurred when reaching the heist state (i.e. if choosing the dark door on trial t-1 had resulted in monetary gain, but the current trial t was a police trial (monetary loss), the consistent choice would be to choose the light door on trial t).

We first conducted this analysis in a separate behavioural experiment (described as “pilot” above; n = 31, see **Methods**). This experiment included exclusively free choice trials, giving us greater power to be able to detect valence effects. In this version of the task, participants were not told about any structure between contexts and integrated information from each context in each condition.

Participants integrated evidence from the other context in the dependent condition (1-4 trials back) in this dataset but also did so in the independent condition - therefore we remain agnostic as to whether participants adjusted how they integrated feedback from state transitions between the two conditions and primarily use this dataset to examine how outcome interacts with learning the state transitions. See **Supplementary Figure 9 and Supplementary Tables 5 and 6** for parameter estimates predicting choice from state transitions experienced 1-5 trials back.] This revealed that transition updating was greater following positive compared to negative outcomes (t(30) = 9.79, p < 0.001, paired sample ttest, **Figure 6a**). In other words, participants updated state transition knowledge and adjusted their subsequent behaviour to a greater degree when outcomes were positive compared to negative. Note that this analysis collapses over contexts (see **Methods**). However the main effect remains (t(30) = 8.14, p < 0.001) when we run this same analysis restricted to the dependent condition.

**Figure 6.**
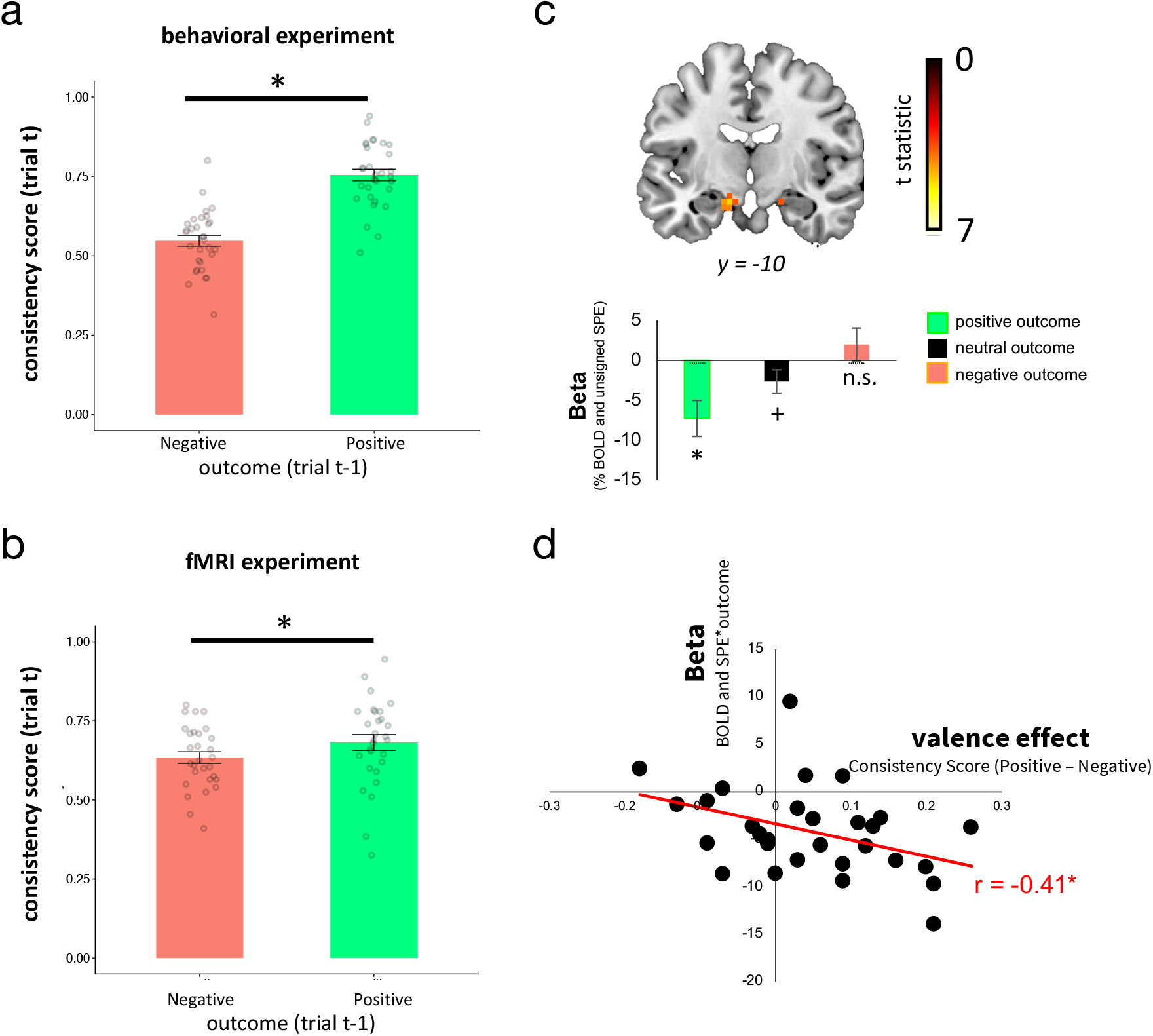
Outcome on the previous trial influenced the degree to which transition knowledge was updated. Specifically, when participants received a positive outcome (+1 on gain trials or 0 on loss trials) consistency scores (indexed as percentage repeat choices for desirable trials and percentage switch choices for undesirable trials) were higher compared to when they received a negative outcome. This was observed in **(a)** participants that completed a behavioural study outside the scanner **(b)** our fMRI cohort. **(c)** Unsigned state prediction errors were modulated by outcome valence in the MTL (peak [x,y,z]: −13, −10, −18), t(28) = 4.87, p = 0.018 FWE whole brain cluster level corrected), image displayed at p < 0.001 uncorrected. **(d)** The magnitude of the valance effect observed behaviourally (quantified as green minus red in (b)) correlated with the size of the interaction betas observed in the fMRI data in (c) (Spearman’s rho = −0.41, p < 0.03). *p < 0.05, ^+^0.05 < p < 0.10: one sample ttest; n.s., non significant

Next, we ran the same analysis on our fMRI participants (restricted to free choice trials, **Figure 6b**). We again observed a main effect of outcome with greater updating following positive compared to negative outcomes (t(28) = 2.24, p = 0.03). This effect remained when analysis was restricted to trials from the dependent condition (t(28) = 2.60, p = 0.015).

In two data sets, our participants’ behaviour suggested that rewards received influenced the degree to which the transition function was updated with a greater update following positive compared to negative outcomes. If this is the case, we would predict that SPE signals in the MTL – which drive updates to the transition function – ought to be larger following positive outcomes compared to negative. To test this, we examined the interaction of the unsigned state prediction error regressor and outcome in a univariate whole-brain analysis (controlling for the main effects of each, see **Methods**). This revealed a negative effect in a cluster in the left MTL (peak [x,y,z]: −13, −10, −18, p = 0.018 whole brain FWE cluster level corrected after thresholding at p < 0.001) which included voxels within our functional ROI (peak [x,y,z]: −16, −7, −18, t(28) = 4.08, small volume corrected using functional ROI mask). No other regions survived whole brain correction. Note that since the main effect of SPE is also negative (**Figure 3a**), the sign of this interaction suggests a greater parametric effect of unsigned SPEs in the MTL following positive versus negative outcomes.

Finally, we examined whether there was a relationship between this interaction effect in the MTL (i.e., the degree to which unsigned SPEs were modulated by outcome valence) and participants’ behaviour (the degree to which consistency scores were greater for positive outcomes compared to negative outcomes). We quantified each participant’s behavioural outcome valence effect (**Figure 6b**) by taking the difference in consistency scores between positive and negative outcomes and correlated these with each participants parametric SPE *outcome interaction beta (this quantifies the degree to which the parametric effect of unsigned SPEs vary are modulated by outcome). This revealed a negative correlation that was robust to outliers (Spearman’s rho = −0.41, p < 0.03); specifically, the greater participants showed a bias towards integrating information following transition sequences that ended in a positive outcome (versus negative) in their choices, the greater extent to which unsigned SPEs expressed in the MTL were greater for higher (versus lower) outcomes.

## Discussion

We studied the neural and computational mechanisms by which humans combine or segment information about the transition structure of the world. We asked participants to learn and track probabilistic state associations in two conditions – one which invited model sharing by integrating information independent of context, and one which invited the use of two separate models, recruited and updated independently, according to the contextual cue displayed. This allowed us to measure differences in behaviour and brain activity as a function of the imperative to either share or keep separate state associations.

We use the term “context” to refer to the visual cues (gem colour) displayed to participants on each trial - a salient feature which participants could use to maintain separate transition functions within a block. Under this definition, participants encountered 2 distinct contexts in both conditions. Participants in our fMRI cohort were explicitly instructed as to which contexts had matching state transition structures. Consequently, these data were best described by a model in which participants switched between compressing these contextual cues (leading to a single task state representation that was context-independent) thereby facilitating model sharing and using these contextual cues (learning two task state representations that were each context-dependent) to maintain separate sets of state transitions. When participants were uninstructed, we observed model sharing in both the dependent and independent conditions. This suggests that *a priori*, participants may have a tendency to believe that items occurring together have shared latent properties such as being associated with correlated transitions, and opt to use a common transition function invariant to sensory cues. This tendency can be overcome in favour of using separate functions with distinct patterns of neural responses for each context via explicit instruction. Our hypothesis is that this would also be the case when this tendency is overcome via learning over time, but this needs to be formally tested.

For the fMRI experiment, we chose to directly instruct our participants, as our hypothesis was agnostic to whether model sharing occurred because of instruction or trial-and-error learning. Consequently, our computational model as currently presented, is an analytic tool and does not offer a process-level account of model sharing. How participants learn and make potentially challenging decisions about which contexts to group together remains a key question therefore. Previous structure learning tasks have suggested that participants are able to use the similarity of latent variables such as value estimates, prediction errors and their covariation over time to draw links between different contexts (Wunderlich et al., 2011, Acuña and Schrater, 2010). Bayesian inference models in which latent causes are inferred and used to group together experiences (Gershman and Niv, 2010, Gershman et al., 2015, Sanders et al., 2020, Niv, 2019) could also be a means that participants learn to group different contexts together in practice. Another possibility is that neural geometry actively represents the relational organisation of task elements (Bernardi et al., 2020, Luyckx et al., 2019, Sheahan et al., 2021). However, our encoding model did not find a smoothly-varying relationship between neural coding and probability, which seems like it would follow naturally from such a representation.

To address the question of how neural population activity (measured at the coarse level of multivoxel patterns) encoded transition probabilities within and between contexts in the dependent and independent condition, we began by identifying voxels that responded differentially according to whether a transition between states was expected or unexpected, using estimates of state prediction errors (SPEs) from the best-fitting (“flexible”) model to parameterise this effect. We found that responses to SPEs correlated with BOLD responses in brain regions that overlapped with the bilateral hippocampus (**Figure 3a**). Of note, the parametric effect of SPEs in the MTL were negative, i.e., this region responded more robustly to expected over unexpected transitions. Whilst this might seem surprising, given the past implication of the (more anterior) hippocampus in novel or surprising stimuli (Strange et al., 2005), in fact the hippocampal-entorhinal BOLD signal has been found to increase to both matches and mismatches in seemingly perplexing ways and often within the same experimental paradigm (Duncan et al., 2012, Henson et al., 2000, Kumaran and Maguire, 2007) and the reasons for enhancement and suppression of MTL BOLD signals by novelty and familiarity remain murky (Barron et al., 2016, Segaert et al., 2013).

We first focussed our analysis on voxels in this region and measured the consistency in neural patterns in these voxels in encoding transition probabilities between conditions. To do this, we needed to account for the fact that participants estimates of these transition probabilities were themselves changing over time, due to fluctuations in the underlying transition dynamics. At some points in the task for instance, choosing the grey door was likely to lead to the heist state, whilst choosing the light door was likely to lead to the neutral state. At other points, the converse was true. One would expect that as these transition dynamics changed, different patterns of BOLD responses would also emerge as encoding adapts to reflect current estimates about the likelihood of going to each state. Therefore, we divided trials into 4 separate quartiles according to participants’ current estimates of transition probabilities inferred from our computational learning model. We then adopted two complementary multivariate approaches: an RSA-based method, which relies on patterns of correlation in fMRI voxels, and a more flexible encoding modelling pipeline, that learned a linear mapping from (model-derived) transition probabilities to fMRI voxels for one session or context and then inverted this model to make predictions in held out sessions or contexts. Both approaches told a similar story: the encoding of probability was more similar across contexts in the dependent condition than in the independent condition. In other words, the brain learned a representation that was similar across contexts when this was beneficial, but partitions probability encoding into different patterns when it is necessary to disambiguate the predictions for different contexts.

The encoding model also enabled us to examine the pattern of results under two different coding schemes – a Gaussian input function and a one-hot input function. Interestingly, whilst the one-hot input function replicated the pattern of RSA and was robust to a range of different probability bins being used, the Gaussian input function did not. We are not entirely clear about why this is the case. Previous theories have emphasised that neural populations in cortex may encode probability distributions in smoothly-varying ways, permitting forms of function approximation or Bayesian inference (Ma et al., 2006, Orhan and Ma, 2017), and there is even some support for this class of theory from studies involving BOLD recordings (Van Bergen et al., 2015). However, the nature of the coding scheme for transition probabilities in hippocampus remains unclear.

The findings described here imply that there is flexibility in the way that transition probabilities are encoded. This potentially allows for “model sharing” – a biological means by which the brain is able to prevent the prohibitive computational cost of planning, by collapsing across contexts with equivalent transition structures. Model sharing may be complementary to other approaches proposed for mitigating the cost of planning. For example, it has been proposed that search paths can be dynamically pruned in ways that focus on the most promising routes (Huys et al., 2012). Other theories, including sophisticated models from AI research, emphasise forms of model compression or abstraction to make planning more effective (Schrittwieser et al., 2020, Wayne et al., 2018). It seems likely that the solution which we propose here – recruiting and updating the same model over different contexts – is one among many ingredients that allow humans to plan effectively.

Observing these effects in the MTL is consistent with past findings that have identified the involvement of the MTL in learning state associations (Eichenbaum et al., 1999, Miyashita, 1988, Yokose et al., 2017, Rey et al., 2018, Schapiro et al., 2012, Schapiro et al., 2013, Deuker et al., 2016, Garvert et al., 2017), encoding relational knowledge that can be used to generalise and draw inferences across contexts (Bunsey and Eichenbaum, 1996, Wimmer and Shohamy, 2012, Zeithamova et al., 2012, Kumaran et al., 2016, Koster et al., 2018, Park et al., 2019) and its role in model based planning (Bradfield et al., 2020), including in two stage sequential planning tasks similar to the one we deploy here (Vikbladh et al., 2019, Miller et al., 2017), potentially via representation of task structure (Geerts et al., 2020). Our initial analysis focused on a region that included different subregions of the MTL. But when we repeated our RSA approach separately in 4 different anatomical subregions of the MTL – hippocampus, entorhinal cortex, amygdala and parahippocampus – we found a significant effect in each of these (an effect which was absent in two control regions). This is suggestive that a network of MTL regions is involved in encoding the predictive relationships between states necessary for planning – consistent with past findings using a similar paradigm to ours (Boorman et al., 2016) – and that each component in this network has the capacity to flexibly adapt the representations it uses to facilitate the sharing of models between contexts when prudent to do so. The involvement of a number of subregions might account for why disabling a specific part of the MTL does not always lead to reductions in goal directed behaviour (Corbit and Balleine, 2000, Gaskin et al., 2005). Interestingly, the effect we observed was strongest in bilateral hippocampus, in line with its involvement in modulating pattern separation between contexts and memories via inputs from other MTL brain regions including the entorhinal cortex (Yassa and Stark, 2011). However, future work – ideally with higher resolution fMRI or direct recordings – is needed to help characterise the precise functional contribution each of these subregions.

We also examined whether there were other regions of the brain in which representations had a similar selective pattern similarity between contexts by running a whole brain searchlight analysis. In addition to confirming the involvement of the MTL, this detected a strong effect in the dorsal striatum and the left IFG. This analysis was exploratory and neither of these brain regions were hypothesised to be involved from the outset. Whilst the IFG and adjacent OFC have previously been shown to be involved in inferring task states using fMRI multivariate approaches (Niv, 2019, Schuck et al., 2016), the striatum was particularly unexpected given its well established role in model free learning (Geerts et al., 2020, Joel et al., 2002, Montague et al., 1996, O’Doherty et al., 2004) although (and with the necessary caveats with regard to retrospective inference), there is some evidence from fMRI and lesion studies that the dorsal striatum – along with prefrontal cortex (Balleine and O’Doherty, 2010, Niv, 2009) – may also play an important role in model based planning behaviour (Yin et al., 2005a, Yin et al., 2005b). Exactly what the functional role either region fulfils here in the service of our task though is unclear.

Computational accounts of model based learning (Sutton and Barto, 1998) typically have the transition matrix stored and updated independently of the reward function. But tasks used to probe choices are not usually designed to test whether this is borne out empirically. For instance, whilst knowledge of the state transitions is used to plan in the Two Step Task – a task often used to measure planning in humans (Daw et al., 2011; Doll et al., 2015) and rodents (Miller et al., 2017) – state transitions are held fixed and do not need to be updated over time. This places all onus on learning the rewards in each state (which gradually change over time). In our task, by having rewards in one state fluctuate between positive (thereby desirable relative to the alternative state) and negative (thereby undesirable relative to the alternative state) and having state transitions change over time we can actually test whether the transition function and the value function are updated independently of one another. This is vital for understanding how planning decisions and deficits thereof (Gillan et al., 2016) are implemented in practice.

Examining participants stay/switch behaviour revealed an effect of valence whereby following positive outcomes, participants updated transition probabilities to a greater degree than following negative outcomes. We note that these findings are unlikely to be accounted for by purely model free state-action learning since our task and updating metric includes cases where participants should (if using model based control and updating the transition function) repeat choices following negative outcomes and switch choices following positive outcomes. These cases would cancel out the effect of valence we actually observe in the data under a model free controller (which would repeat following positive and switch following negative outcomes). An effect of valence on updating was also observed in the fMRI data which revealed a greater parametric effect of SPEs for positive outcomes relative to negative in the MTL.

Normative accounts of planning dissociate representations of the value function and the transition function so that agents can learn about the world independent of specific outcomes. So why might – as our data suggests – humans have evolved to modulate learning in such a way that deviates from these normative accounts? Sensitivity to new information can vary according to many factors such as surprise (Pearce and Hall, 1980), volatility (Behrens et al., 2007) and arousal levels (Dundon et al., 2020, Garrett et al., 2018, Li et al., 2011). One possibility is that the valence of the outcome received acts as a cue which modulates learning (via arousal for instance). Interestingly, the pattern of asymmetric updating is reminiscent of confirmation bias (Nickerson, 1998) whereby learning is greater when information confirms prior beliefs. In bandit tasks this manifests itself as greater learning for positive outcomes from bandits that were chosen and negative outcomes for unchosen bandits compared to learning from negative outcomes for chosen bandits and positive outcomes for unchosen bandits (Fontanesi et al., 2019, Palminteri et al., 2017, Tarantola et al., 2021). A recent account of confirmation bias (Lefebvre et al., 2020) has used reinforcement learning models to show that this learning asymmetry can in fact be beneficial by driving apart the difference in value between the different options, thereby helping agents choose the correct option in the face of decision noise. Future theoretical work may help shed light on whether a similar normative account exists behind the asymmetry we observe here in planning to show that that biasing learning towards pathways that lead to reward (rather than avoid punishment) could help maximise reward in model based planning under certain conditions.

Together these results shed important light on the computational processes by which the MTL maintains and adapts knowledge about the consequences of our choices and actions in the world. First, they demonstrate that by relying on a common representational code, knowledge can be shared across different contexts that we interact with. This is essential for learning in the natural world where the sheer number of contexts we deal with makes maintaining exclusively context specific representations unrealistic whilst at the same time inappropriate model sharing (e.g., between contexts with distinct transition structures) is detrimental. Second, by modulating learning via outcomes, state associations can be revised faster following rewards – this could be beneficial for helping us strengthen associations that lead to positive outcomes and forget those that don’t but may be maladaptive when states and rewards fluctuate quickly and independently of one another.

## Methods

### Participants

A total of 62 healthy volunteers with no self-declared history of psychiatric or neurological disorders took part in the experiment. Thirty one took part in the pilot experiment (18 female, mean [std.] age: 26.29 [5.50]), and thirty-one participated in the main fMRI study. From the latter, two participants were subsequently excluded. One was excluded because their structural fMRI revealed a possible brain abnormality. A second participant was excluded due to excessive head motion (more than 10% of images contained motion artefacts upon visual inspection). This left 29 participants (16 female; mean [std.] age: 25.86 [3.59]) in the final sample. Participants were paid 10€/hour plus a bonus contingent on performance.

### Ethics Statement

The fMRI study was approved by the ethics committee of the University of Granada where data collection was carried out. All participants gave written informed consent prior to scanning. The behavioural pilot was approved by the ethics committee at the University of Oxford where this data set was collected. We obtained written informed consent from each participant.

### Heist Task

On each trial of the fMRI experiment, participants were presented with one of two doors (dark/light) on one (left/right) side of the screen (side counterbalanced), in one of two “contexts” within the current block (**Figure 1a**). On forced trials (24 per trials per block), participants were required to select the door initially presented. On free choice trials (8 per block), participants could either choose the door initially presented or opt to choose the alternate door which appeared on the opposite side of the screen (**Figure 1a**). Participants had 1.5 seconds to respond otherwise the trial aborted. Missed trials (mean [std] = 6.41 [4.65]) were excluded from all analyses.

The selection of door influenced which of two possible 2^nd^ stage states participants subsequently transitioned to. One of the doors transitioned with probability *p* to a heist state where participants could either win or lose money and transitioned with probability 1 - *p* to a neutral state in which participants would always receive 0 as an outcome (participants were only rewarded for free choice trials). The alternate door transitioned to the same second stage states but with the inverse probability (i.e., probability 1-*p* of transitioning to the heist state and probability *p* to the neutral state, **Figure 1b**). The value of *p* was was set to either 0.2 or 0.8, alternating randomly between these two values throughout the task (with probability of changing equal to 0.1 on every trial). This meant that one door was always likely to transition to one of the outcome states and unlikely to transition to the other. Participants were told state transitions could change but were not told the probability with which this could happen. Importantly, *p* always had the same value for both contexts in dependent blocks. In independent blocks, the values for *p* were independent in the two contexts. Participants were explicitly told this probability structure during the instructions and the block type they were in (dependent/independent) was clearly signalled to them at the start of a new block of trials. The context of the current trial was signalled to participants by the colour of a gemstone presented in the centre of the screen (either green, yellow, blue or red). The assignment of gemstone to context was different for each participant but (after assignment) remained the same throughout the experiment. Whether a gain or a loss was possible in the heist state alternated on each trial and was signalled to participants during door and response presentation. Explicitly providing participants with this information was done to remove the need to actively learn the value of each bottom level state and emphasise the need to track the transition function and use current beliefs about this function to plan. After indicating their choice, participants were shown the state they transitioned to and the resulting outcome – either a gain or a loss if they transitioned to the heist state or zero if they transitioned to the neutral state.

The task took place in sessions of trials (2 blocks of 32 trials per session, 5 sessions total during the experiment, 320 trials total). The first session took place outside of the scanner. Each session contained one block of trials in the dependent condition and one block of trials in the independent condition. The order of the blocks was counterbalanced across sessions. Participants indicated their response using a computer keyboard (outside the scanner) or MRI compatible button box (inside the scanner). Participants were paid a base-rate bonus of 2.50 Euro plus 2.5 times their percentage of correct free choice trials (up to 5 Euro total). The task was programmed in MATLAB using Psychtoolbox (Kleiner et al., 2007).

### Behavioural analysis (adapting information integration between contexts)

To examine the extent to which participants updated beliefs about state transitions within and between contexts, logistic regression analyses were conducted (mixed-effects models using the fitglme fitting routine in MATLAB, version 2020 [https://www.mathworks.com/]). Models tested to what extent subjects’ choice behaviour on each trial (coded as: select dark door = 1; select light door = 0) was influenced by transitions experienced over the previous 5 trials.

To examine this, we first constructed 5 variables that coded the evidence received from the state transition n trials back (relative to the current trial, t), where n ranged from 1 to 5. When trial t was a gain trial, previous transitions to the heist state were coded 1 (−1) if the dark (light) door was selected t-n trials back and participants transitioned to the heist state and coded −1 (1) if the transition encountered was to the neutral state. This coding was reversed for loss trials (**Figure 2**). The intuition implicit in this coding scheme is that participants would aim to repeat choices that previously transitioned to the heist state on gain trials but to switch choices on loss trials (in an attempt to transition to the neutral state and avoid incurring a loss). We also partitioned trials according to whether evidence was received in the same or alternate context as the current trial t. This lead to a total of 10 variables – 5 encoding evidence received 1 to 5 trials back from the same context and 5 encoding evidence received 1 to 5 trials back from the alternate context. 0 was entered as a value for cases where a variable did not apply for a particular trial (for example, if 3 trials back a subject’s choice was executed in the alternate context, evidence 3 trials back in the same context would be assigned a value of 0 for this trial).

Next, to assess *qualitatively* whether the degree of information integration from each context (same and other) changed between conditions, we entered all 10 variables in separate mixed effects models: one for the dependent condition and one for the independent condition. Only choices from free choice trials were entered in the model as the dependent variable (however the information encoded in the independent variables used to predict choice could come from free or forced trials as participants could use transition information from both trial types). All regressors and the intercept were taken as random effects, i.e. allowed to vary across subjects.

The model was specified in the syntax of MATLABs fitglme routine as:

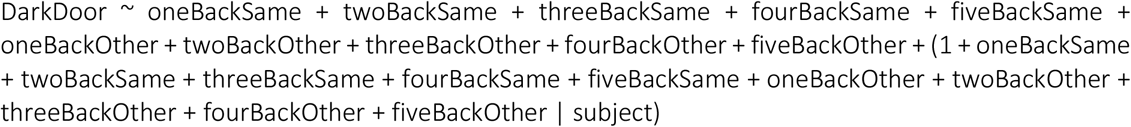

To the extent that participants are using information from each context to a similar degree (which ought to be the case in dependent blocks), coefficient estimates ought to have a similar magnitude for same and other context. To the extent that participants ignore information from an alternate context to a similar degree (which ought to be the case in independent blocks), there ought to be separation between coefficient estimates from same versus other. Note that by controlling multiple trials back we guard against the possibility that information used in the alternate context can have an effect in the dependent condition by virtue of the fact the feedback of received is similar in the two contexts.

Finally, to assess *quantitively* whether differences in information integration between conditions were significant, we averaged each condition’s streams of evidence for picking the dark door on the current trial over the past five trials. This resulted in two quantities:

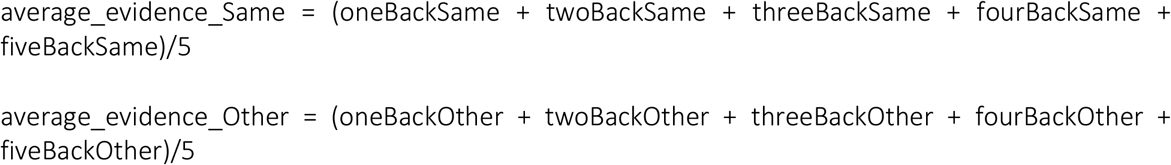

We then subtracted average_evidence_Other from average_evidence_Same providing a difference score:

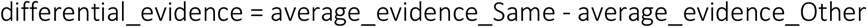

The differential evidence score reflects a relative preference in updating beliefs for information received from the same context over information received from the other context. When equal to 0, individuals are indifferent between evidence from the same and evidence from the other context. When greater than 0, individuals prefer (i.e. update beliefs to a greater degree) information received in the same context compared to the other context. When less than 0, individuals prefer information received in the other context compared to the same context.

We used this differential evidence score in a third mixed effects model to test whether preferences for the context in which information was received shifted with condition (captured in the model as a Differential Evidence by Condition interaction). The model was specified as follows:

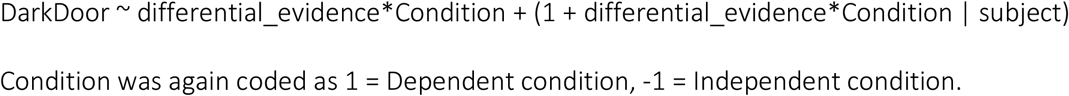

### Computational Model

Our model is not intended primarily as an account of the computations that humans undertake, but as an analytic tool. Participants are assumed to track the task’s underlying state transition structure in the form of *p*, an estimate of the probability that selection of one of the two doors (which of the two is arbitrary, but in our modelling this is taken to be the dark door) transitions to the heist state. This is assumed (as is the actual case in the experimental design) to be equal to the probability that the alternate door transitions to the neutral state. It is also assumed (as is the case) that 1-*p* is equal to the probability of each door going to the alternate state (dark goes to neutral and light to heist). Under these assumptions, maintaining a belief about a single quantity, *p*, enables computation of estimates for each door going to each 2^nd^ level (terminating) state. Importantly, participants are assumed to maintain two sets of beliefs about *p*: 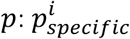 and *p*_*independent*_. 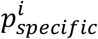, maintains separate estimates of *p*, exclusive to each context where i indexes the 2 contexts in each block (i.e. 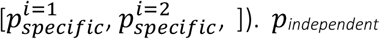 maintains a single estimate of *p* which updates across contexts (within the same block). All estimates of *p* were initialised to 0.5 at the start of the experiment in all models.

At the time of choice, participants then combine the two sets of beliefs 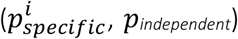 into a single estimate, 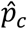, according to:

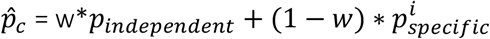

We tested a baseline model in which *w* was held fixed between conditions. We refer to this as the *fixed model*. We tested this against a 2^nd^ model which was identical in all respects except that it allowed *w* to reverse in the independent condition. In other words, in the dependent condition 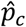 was calculated as:

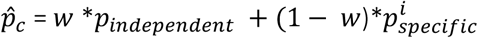

In the independent condition 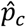was calculated as:

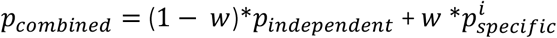

We refer to this as the *flexible model*.

In both models, combined estimates of 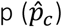 were then used to calculate the value of selecting each

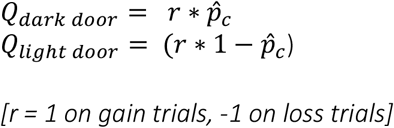

Following choice, after participants observed the 2^nd^ level state they transitioned to, a state prediction error, δ, calculated as:

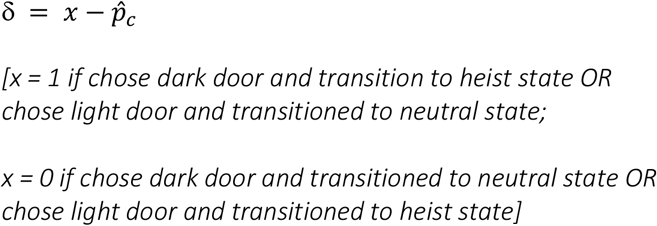

This prediction error was then applied to update both sets of beliefs about p:

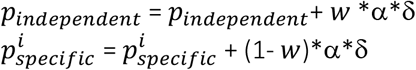

Where the context indexing 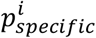^*i*^ can be context 1 or context 2.

The w used in each update is identical to w used to compute 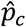 and was either held fixed (fixed model) or allowed to reverse between conditions (flexible model).

To avoid probability estimates exceeding 1 or going below 0 (which in a small number of cases is possible in this setup), updates to beliefs were bounded to within this range.

The probability of choosing the dark door was then estimated using a softmax choice rule, as follows:

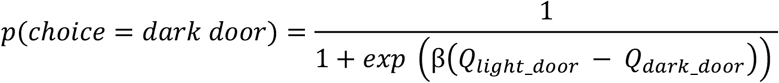

Altogether each model has 3 parameters: α, β and *w*. For each participant, we estimated the free parameters of the model by maximizing the likelihood of their sequence of choices, jointly with group-level distributions over the entire population using an Expectation Maximization (EM) procedure (Garrett and Daw, 2020, Huys et al., 2011) implemented in the Julia language (Bezanson et al., 2012), version 0.7.0. Note, similar to the behavioural analysis reported above, all trials (forced and free) were included in the model but only free choice trials were included in the likelihood calculation. Models were compared by first computing unbiased per subject marginal likelihoods via subject-level cross-validation and then comparing these likelihoods between models (Flexible versus Fixed) using paired sample t-tests (two sided).

### Computational Simulations

To examine the qualitative fit of each learning model to the data we ran separate simulations for the Fixed Model (in which w was held constant across conditions) and the Flexible Model (in which w was allowed to vary with condition). For each simulation (n = 504 for each model), we ran a group of 29 virtual participants. For each virtual participant, we randomly selected (with replacement) a set of parameters (*β, α* and w) from the best fit parameters generated by the computational model (fit to actual participants choices). We then simulated the learning process by which estimates of p evolved (given door selection and state encountered), exactly as described for the respective computational models. To mimic the task as closely as possible, 25% of a virtual agents trials were free choice trials in which we simulated which of the two doors were selected (given current beliefs about p, and whether a gain or a loss was available in the heist state) and 75% were forced choice trials where the door selected was chosen for them (as a coin flip).

We then entered choices made by each virtual agent as the dependent variable in a binomial mixed effects model with regressors coding evidence received 1 to 5 trials back from the same and alternate context (10 regressors in total). This was run separately for each condition replicating the analysis conducted on the data (i.e. actual subjects’ choices) with the same model specification (as before, all regressors and the intercept were taken as random effects). This generated a set of fixed effect parameter estimates for each simulation for each condition. We then averaged each fixed parameter estimate over the simulations and compared these to the parameter estimates generated from the data.

Finally we used the fixed model to run a permutation test to estimate the extent to which an interaction between differential evidence and condition (our third mixed effects model) could arise under agents that did not change information integration between contexts which might occur due to feedback being more similar in the dependent condition compared to the independent condition. Specifically, we simulated choices for 500 groups made up of 29 agents each, performing the task. For each agent, we randomly selected (with replacement) a set of parameters (*β, α*) from the best fit parameters generated by the fixed model (fit to actual participants choices). ***W*** could take any value between 0 and 1 (uniformly distributed) and could not reverse between contexts. For each group we then calculated differential evidence scores on each trial for each participant and entered these into a mixed effects model to predict choices (along with condition and their interaction) exactly as we did using participants data. This generated a distribution of fixed effects estimates and t statistics which we used to calculate a 95% confidence interval and compare against the estimates found in the data.

### fMRI image acquisition, pre-processing and reporting

MRI data were acquired on a 3T Siemens Magnetom Trio MRI Scanner (Erlangen, Germany) scanner. A whole brain high-resolution T1-weighted anatomical structural scan was collected before participants commenced the four in-scanner blocks of the task (imaging parameters: 1mm^3^ voxel resolution, TR = 1900ms, TE = 2.52ms, inversion time (TI) = 900ms, slice thickness = 1mm, voxel resolution = 1mm^3^). During the task, axial echo planar functional images with BOLD-sensitive contrast were acquired in descending sequence (imaging parameters: 32 axial slices per image; voxel size = 3.5mm^3^, slice spacing = 4.2mm, TR = 2000ms, flip angle = 80°, TE = 30ms). 462 volumes were collected per participant per session (total number of volumes over the 4 sessions = 1848), resulting in a scanning time of approximately 1 hour. Image analysis was performed using SPM12 (http://www.fil.ion.ucl.ac.uk/spm). The following procedures were used for preprocessing of the raw functional files. Slice-time correction referencing was applied with reference to the middle slice to correct for/avoid interpolation errors due to the descending image acquisition sequence (Sladky et al., 2011 in Juechems et al. (2017)). Then, realignment of thet images from each session with the first image within it was performed. The crosshair was adjusted to the anterior commissure manually to improve coregistration. After coregistration of the functional with the structural images was performed, segmentation, normalisation and smoothing of the epi files was undertaken. We then checked for motion artefacts and flagged scans as well as warping manually.

In all fMRI analysis (univariate and RSA searchlights) we report activation that survives small volume correction at peak level within an anatomical or functional ROI mask (see below for how these were defined). Other brain regions were only considered significant at a level of p < 0.001 uncorrected if they survived whole-brain FWE correction at the cluster level (p < 0.05).

### Anatomical masks

Anatomical masks were generated using the automated anatomical labelling (AAL) atlas (Tzourio-Mazoyer et al., 2002) and Talairach Daemon Atlas (Lancaster et al., 2000), which was used to define Brodmann area 28 as entorhinal cortex (Canto et al., 2008) and Brodmann area 17 as V1 (Tootell et al., 1998) integrated in the WFU Pickatlas GUI (Maldjian et al., 2003):

1. A bilateral medial temporal lobe mask used for small-volume correction, which was defined as including the bilateral hippocampus, entorhinal cortex, parahippocampus and amygdala and dilated by a factor of 1 in the WFU Pickatlas GUI.
2. Bilateral amygdala (84 voxels), hippocampus (336 voxels), entorhinal cortex (53 voxels), and parahippocampus (404 voxels) masks (no dilation) used for anatomical definition of our ROI from fMRI general linear model 1 (see below) as well as post-hoc RSA tests (**Figure 4 & Supplementary Figure 6**).
3. Bilateral V1 (121 voxels) and bilateral primary motor cortex (Brodmann area 4, 240 voxels) used as a control region for the post-hoc RSA tests (see **Figure 4** & **Supplementary Figure 6**).

All masks were resliced to match the dimensions of our data using the SPM fMRI Realign (Reslice) function.

### fMRI general linear model 1

For each participant, the blood oxygen level-dependent (BOLD) signal was modelled using a General Linear Model (GLM) with time of door presentation and time of outcome presentation as onsets. Events were modelled as delta (stick) functions (i.e. duration set to 0 seconds) and collapsed over our two experimental conditions (dependent and independent blocks).

To identify regions tracking state prediction errors, we extracted trial by trial estimates of unsigned state prediction errors, |δ|, from our computational model and entered these as parametric regressors, modulating the time of outcome for each participant. In addition, we also entered the following regressors: outcome received (1, 0 or −1), the interaction of outcome with unsigned state prediction error (i.e. the product of outcome received with |δ| on each trial) and trial type (1 = forced, −1 = free). Six movement parameters, estimated from the realignment procedure were added as regressors of no interest.

### ROI definition

We identified region(s) in which the BOLD response was parametrically modulated by the magnitude of the unsigned state prediction error (|δ|), using a threshold of p < 0.001 uncorrected, with cluster size > 10 voxels. Clusters identified were saved as binary regions of interest (ROIs; in SPM) and then combined into a single ROI using the MarsBaR toolbox (http://marsbar.sourceforge.net/). This functional ROI was then used for subsequent representational similarity analysis (RSA; see below). We divided the number of voxels that fell within both our functional ROI and each anatomical mask by the total number of voxels in our functional ROI. This gave us the percentage with which our functional ROI was a conjunction of each anatomical region.

### fMRI GLM 2a (door presentation)

For each participant, we created a design matrix in which each door presentation (32 per condition per session) was modelled as a separate event (without parametric regressors attached). Such a procedure has been used multiple times in the past (Charpentier et al., 2014, Garrett et al., 2016). Outcome onset was entered as an additional event. Events were modelled as delta functions and convolved with a canonical hemodynamic response function to create regressors of interest. Six motion correction regressors estimated from the realignment procedure were entered as covariates of no interest.

### RSA (door presentation)

To examine whether BOLD responses were more similar between contexts in the dependent versus independent condition, we used GLM3a to extract estimates of BOLD response on each trial in our functional ROI (identified from GLM 1) and partitioned these estimates into four linearly spaced bins according to how likely the door presented was to go to the heist state (P(state = heist | door_presented)). This was inferred by extracting trial by trial estimate of *p*_*combined*_ (from the flexible learning computational model) and using *p*_*combined*_ or 1-*p*_*combined*_ depending whether the dark or light door was presented respectively to estimate *p*(state=heist | door_presented).

We divided trials into quartiles based on *p*(heist state | door presented), resulting in the following average (standard deviation; SD) probability bins:

Bin 1: 0.04 < *p* (heist state | door presented) ≤ 0.21 (0.10)

Bin 2: 0.21 < *p* (heist state | door presented) ≤ 0.51 (0.09)

Bin 3: 0.51 < *p* (heist state | door presented) ≤ 0.80 (0.09)

Bin 4: 0.80 < *p* (heist state | door presented) > 0.96 (0.03)

This was done separately for each context that the participant (N = 29) encountered (2 in dependent blocks and 2 in independent blocks, 16 bins in total). We then averaged these estimates in each voxel in our functional ROI (collapsing across the 4 functional runs) for each bin generating an average BOLD response for each voxel.

To compare the similarity of responses between contexts we proceeded by first calculating the dissimilarity of BOLD responses in each of the 4 bins between contexts. This was computed using the pdist function in MATLAB using 1-pearson correlation as a measure of distance; hence high correlation indicates a low level of dissimilarity (conversely a high level of similarity). This generated an 8×8 dissimilarity matrix for each condition of which we subselected the 4×4 matrix displaying the dissimilarity of probability bins between the two contexts (i.e. context 1 vs context 2 for each level of *p* (heist state | door presented))

Dissimilarity scores were then converted into similarity scores (high scores indicating greater similarity) and Fisher transformed to allow inference at the group level. The four similarity scores along the diagonal of each RSA matrix (where identical bins are compared between contexts) were averaged for each participant creating an on-diagonal similarity score which quantifies the extent to which identical values of transition probabilities are encoded similarly between the two contexts in a condition. The 12 similarity scores on the off-diagonal of each RSA matrix (where different bins are compared between contexts) were separately averaged together to create off-diagonal similarity scores. Note that unlike in regular RSA analyses, all 12 scores were averaged across rather than just the upper or lower triangle as the values in the 4×4 RSM are not identical about the diagonal (off-diagonal 4×4 of a larger 8×8, see above). We then computed the difference between on and off diagonal scores separately for each condition. One sample ttests (versus 0) were conducted to assess whether significant differences between on an off diagonal similarity scores existed. Two-tailed paired sample ttests were used to compare whether difference scores were greater for the dependent condition compared to the independent condition. For a crossvalidated version of this analysis in which where we exclude within block autocorrelations, see **Supplementary Figure 3**.

The same RSA procedure was applied to voxels within the four anatomical ROIs used to characterise the nature of the effect within the medial temporal lobe and the control regions V1 and M1 (**see Figure 4**). The interaction ANOVA result reported in-text are Greenhouse-Geisser corrected to adjust for violations of sphericity (both F value and degrees of freedom).

To check whether there is a relationship between the temporal proximity of trials between contexts and how similar the neural patterns are, we calculated the mean temporal distance between trials in the two contexts on the diagonal and the off-diagonal in each condition for each participant. We then correlated the difference in proximity between diagonal and off-diagonal trials with the difference in representational similarity between the diagonal and off-diagonal in the dependent and the independent condition (scatter plots of these correlations are presented in the **Supplementary Figure 8**).

### Encoding model analysis

As a complementary approach, we built a linear encoding model, equivalent to a crossvalidated multinomial logistic regression, that mapped voxels (within an ROI) onto probabilities under different constraints. We evaluated this model in cross-validation, using independent held out data from across scanner runs. Briefly, we first extracted single-trial estimates of BOLD within the MTL ROI for each gem on each session, yielding data *Y* of size *v* × *t*, where *v* is the number of voxels and *t* the number of trials on which that gem was presented. We also recoded (scalar) single-trial, model-derived estimates of transition probability (converted to odds ratios) as input vectors in either a one-hot format (i.e. a one within the relevant bin and zeros elsewhere) or a Gaussian format (i.e. a Gaussian tuning curve that was maximal in the relevant bin but gradually tapered over adjacent bins). We used *n* bins falling within the range (in log odds) units of −2 to 2, where *n* varied exhaustively from 1-10. This yielded data *X* of size *n* × *t*. We estimated weights *w* by linear regression of *X*_*i*_ onto *Y*_*i*_ for scanner run *i* and evaluated the fit of the model to help out probabilities *X*_*j*_ from multivariate patterns *Y*_(_ acquired in scanner run *j*. We used a (mean) cross-entropy loss in validation. This exercise allowed us to verify, for each gem, the cross-validated loss when weights obtained with gem *g* were evaluated with gem *g*′ with which it co-occurred, both in the independent condition (where the probabilities were different) and the dependent condition (where they were not). We tested whether there was stronger cross-validation between gems (and across runs) in the dependent than the independent condition, for varying number of bins *n* and with both one-hot and Gaussian input functions.

### Searchlight RSA analysis (door presentation; whole-brain)

To assess whether our ROI was the only brain area with dependent and independent block transition probability representations and potential differences between them or whether this representation was distributed across the brain (and thus potentially less meaningful), we also conducted a whole-brain searchlight analysis. The searchlight analysis was conducted using a combination of scripts from the RSA toolbox (Nili et al., 2014) and our own parser script feeding in the single-trial onset events generated in GLM3a. The searchlight radius used was 10.5mm (corresponding to 3 voxels). Neural representational dissimilarity maps for the two block types were separately correlated with model representational dissimilarity matrices (RDMs, **Figure 3d**) using Spearman’s correlation coefficient. The model RDM specified that the on-diagonal was more similar between contexts than the off-diagonal. This was done individually for each participant and the resulting maps of correlation coefficients were saved. Second-level analysis as described above was then applied to the r-maps to establish separate group-level effects for the two conditions, i.e. the dependent and independent blocks (**Figure 3e**). We report any brain regions that survive whole brain correction at the cluster level after thresholding at p < 0.001.

### fMRI GLM 2b (outcome presentation)

For each participant, we created a design matrix in which each outcome presentation (32 per condition per session) was modelled as a separate event (without parametric regressors attached). Door presentation onset was entered as an additional event.

### RSA analysis (outcome presentation)

We used GLM2b to extract estimates of BOLD response on each trial in our functional ROI and partitioned these estimates into bins according to the combination of doors chosen and state encountered. These combinations (of which there are 4 in total) drive the direction and degree of update of beliefs (p) about state transitions in the computational model. Specifically, we divided responses into bins as follows:

Bin 1: dark door chosen + heist state encountered

Bin 2: dark door chosen + neutral state encountered

Bin 3: light door chosen + heist state encountered

Bin 4: light door chosen + neutral state encountered

This was done separately for each context that the participant encountered (2 in dependent blocks and 2 in independent blocks, 16 bins in total). We then averaged these estimates in each voxel in our functional ROI (collapsing across the 4 functional runs) for each bin generating an average BOLD response for each voxel.

To compare the similarity of responses between contexts we followed a similar procedure to the RSA analysis conducted at door presentation. We first calculated the dissimilarity of BOLD responses in each of the 4 choice-outcome state combinations across the two conditions generating 2 separate 8*8 RSA matrices of which we subselected the off-diagonal 4×4 for further analyses (context 1 vs context 2 for each of the four choice-outcome state combinations computed separately for each condition). After conversion to similarity scores and Fisher transformation, the four on-diagonal similarity scores and the 12 off-diagonal similarity scores of each RSA matrix were averaged to create 2 sets of similarity scores per condition. The mean on- and off-diagonal similarity scores were then entered into a paired ttest to assess differences between identical choice-outcome bins and non-identical choice-outcome bins in the two contexts. Then, to assess whether there were meaningful differences between conditions, the difference between the mean on and off-diagonal scores for each participant in each condition was entered into a paired ttest (dependent on-diagonal-off-diagonal vs independent on-diagonal vs of-diagonal).

The same RSA procedure was applied to voxels within the four anatomical ROIs used to characterise the nature of the effect within the medial temporal lobe and the control regions V1 and M1 (see **Figure 5**). Again, the ANOVA results reported were Greenhouse-Geisser corrected due to violations of the assumption of sphericity.

### Searchlight RSA analysis (outcome presentation; whole-brain)

The searchlight analysis was implemented in the same way as described above for the searchlight RSA at time of door onset. Here, the onset events read into the searchlight script were the outcome onset events generated in GLM3b. Again, the model RDMs specified that the on-diagonal (identical choice-outcome combinations for the two contexts within a condition) was more similar than the off-diagonal (**Figure 5a**) and the analysis was conducted separately for the two conditions.

### Searchlight interaction analysis (outcome presentation; whole-brain)

The interaction analysis was also conducted similarly to the analysis described above for the time of door onset. In this case, if there is a difference between the difference scores for the two conditions, this means that the difference between the similarity in encoding of identical choice-outcome combinations and different choice-outcome combinations across the two contexts is different between the two conditions. If this difference is positive (as this analysis is coded as encoding similarity), it means the same choice-outcome combinations are encoded more similarly between contexts than non-identical choice-outcome combinations in dependent than in independent blocks and vice versa. As for door presentation, we report any brain regions that survive whole brain correction at the cluster level after thresholding at p < 0.001.

### fMRI general linear model 3

To visualise the parametric effect of our interaction term (|δ| * outcome) in GLM1, we ran a separate GLM which included onsets of door presentation and outcome presentation with outcome onsets separated into 3 separate events: time of outcome presentation when participants received an outcome of +1, an outcome of 0 and an outcome of −1. Each of the 3 outcome onsets was modulated by 2 parametric regressors: unsigned state prediction error (extracted from our flexible RL model) and trial type (force/free). Events were modelled as delta functions and collapsed over our two experimental conditions (dependent and independent blocks), just as for fMRI GLM1. Six movement parameters, estimated from the realignment procedure were added as regressors of no interest. We then extracted the parametric betas for the state prediction error regressors for each participant from the 3 outcome conditions using the Marsbar toolbox at the peak voxel of the |δ| * outcome cluster identified in GLM1.

### Participants and task (behavioural pilot)

Thirty-one self-declared healthy individuals (18 female; *M* = 26.29 years, *SD* = 5.50) were recruited using opportunity sampling via the Oxford University Research Recruitment System. The task was the same as the fMRI cohort undertook described above) save for the following differences. Firstly, participants performed 8 blocks of 60 trials (480 trials total) and all trials in this design were free choice trials. This provided us with a higher powered design to detect differences in updating due to outcome received at the end of an episode. After an inter trial interval (0.3-0.5s) participants had up to 5 seconds to make their choice after which they received confirmation of their choice (0.5s) and feedback (1s). Second, participants were not informed about the differences between blocks. However, just as before each block had two different contexts, a dependent block in which transitions for the two contexts was the same and an independent block in which the transitions were independent.

### Behavioural analysis (outcome valence and state transition updating)

To examine the effect of outcome valence on transition updating we calculated a consistency score for each participant. This is the percentage of times a participant’s choices were consistent given both: (1) the previous trials state-action-state sequence, (2) whether the current trial was a gain or a loss trial. Since the same state action state sequence can lead to repeating or switching being the correct thing to do - depending whether the next trial is a gain or a loss trial – we first divided trials into two types - repeat and switch. Repeat trials are those for which participants would want to revisit the terminating state from the previous trial. For example, participants would want to repeat their choice if they picked the grey door on the last trial, went to the vault and the next trial is a gain trial. These trials comprised:

i. Trials where they previously reached the heist state AND the current trial was a gain trial.
ii. Trials where they previously reached the neutral state AND the current trial was a loss trial.

*Switch trials* are those where participants would want to avoid the terminating state from the previous trial. For example, participants should want to swich their choice if they picked the grey door on the last trial, went to the vault and the next trial is a loss trial. These trials comprised:

i. Trials where they previously reached the heist AND the current trial was a loss trial.
ii. Trials where they previously reached the neutral AND the current trial was a gain trial.

For both repeat and switch trials, the outcome on the previous trial can be positive or negative. For instance, whilst a participant ought to want to repeat selection of grey door if that took them to the vault on the last trial and the next trial is a gain trial, the outcome on the last trial (when they went to the vault) could have been positive or negative, depending whether the last trial was a gain or a loss trial. Hence we then further divided each trial type – repeat, switch – into those where they received a positive (+1 on gain trials, 0 on loss trials) or negative (−1 on gain trials, 0 on loss trials) outcome at the end of the previous transition. This gave us 4 types of trials – repeat positive, repeat negative, switch positive and switch negative. We calculated the % of trials participants repeated or switched choices (as appropriate) for these 4 trial types for each participant. We then calculated a *consistency score* for positive trials by averaging together repeat positive and switch positive. We also did the same for negative trials.

For the behavioural experiment dataset, all trials were used. In the fMRI dataset, only free choice trials were included (but transition sequences from the previous trial could be from a free or a force trial). Participants’ consistency scores for positive were compared to negative using paired sample ttests (two tailed). First we did this collapsing over contexts and conditions. This meant that the previous trial could have either been from the same or from the alternate context. Note that participants were not explicitly told of the conditions (i.e., whether to ignore or take notice of contextual cues) in the behavioural dataset. Although they were told this in the fMRI version of the task this ought not to bias this analysis. Nonetheless, we also repeated this analysis only using trials in the dependent condition.

Finally, we calculated a quantified each participants outcome valence effect as the difference between consistency scores for positive trials (i.e. repeat positive and switch positive trials) minus consistency scores for negative trials (i.e. repeat negative and switch negative trials). This indexed the degree to which participants updated state transitions preferentially following positive compared to negative outcomes over both types of trials. We then correlated each participants valence effect with their parametric betas extracted for the interaction regressor (|δ| * outcome) from GLM1.

## Acknowledgements

We thank Tania Martinez Montero and Alberto Sobrado for assistance with fMRI scanning. We would like to thank Dan Bang for helpful insight and comments on an earlier draft of the manuscript and Nathaniel Daw for the EM code used for model fitting and comparison.

## Financial Disclosure Statement

This research was funded in part by the Wellcome Turst (Sir Henry Wellcome Postdoctoral Fellowship to N.G., grant reference: 209108/Z/17/Z), a European Research Council Consolidator Grant to C.S. as well as support from the Human Brain Project (Special Grant Agreement 3) to C.S. and a Waverley Scholarship to L.G.. The funders had no role in study design, data collection and analysis, decision to publish, or preparation of the manuscript. For the purpose of open access, N.G. has applied a CC BY public copyright licence to any Author Accepted Manuscript version arising from this submission.

## Competing interests

The authors declare no competing interests.

## Data Availability

Behavioural data and analysis scripts for all analyses are available at: https://github.com/summerfieldlab/Garrett_Glitz_etal.fMRI data (2^nd^ level SPM maps and similarity scores in regions of interest) are available at: https://osf.io/zvkj3/

## Supplementary Material

**Supplementary Table 1.** Dependent Condition: Fixed effect coefficients from Lagged Logistic Regression Model (N=29).*p<0.05, **p<0.01, ***, p<0.001

| Name | Estimate | SE | tstat | pValue | Lower | Upper |  |
| --- | --- | --- | --- | --- | --- | --- | --- |
| Intercept | -0.12234 | 0.098273 | -1.2449 | 0.21346 | -0.31518 | 0.0705 |  |
| oneBackSameGem | 1.1124 | 0.17782 | 6.2556 | 5.80E-10 | 0.76345 | 1.4613 | *** |
| twoBackSameGem | 0.43516 | 0.13316 | 3.2679 | 0.0011192 | 0.17386 | 0.69646 | ** |
| threeBackSameGem | 0.12169 | 0.13769 | 0.88379 | 0.37702 | -0.1485 | 0.39188 |  |
| fourBackSameGem | 0.12764 | 0.14018 | 0.91055 | 0.36274 | -0.14743 | 0.40272 |  |
| fiveBackSameGem | 0.27492 | 0.13963 | 1.9689 | 0.049235 | 0.0009229 | 0.54892 | * |
| oneBackOtherGem | 0.71938 | 0.15133 | 4.7537 | 2.28E-06 | 0.42243 | 1.0163 | *** |
| twoBackOtherGem | 0.40005 | 0.11724 | 3.4122 | 0.0006696 | 0.16999 | 0.63011 | *** |
| threeBackOtherGem | 0.045014 | 0.12806 | 0.35151 | 0.72528 | -0.20627 | 0.2963 |  |
| fourBackOtherGem | 0.054181 | 0.12789 | 0.42364 | 0.67191 | -0.19678 | 0.30514 |  |
| fiveBackOtherGem | 0.088797 | 0.12254 | 0.72461 | 0.46885 | -0.15167 | 0.32926 |  |

Fixed effects coefficients

**Supplementary Table 2.** Independent Condition: Fixed effect coefficients from Lagged Logistic Regression Model (N=29).*p<0.05, **p<0.01, ***, p<0.001

| Name | Estimate | SE | tstat | pValue | (95% CIs): |  |  |
| --- | --- | --- | --- | --- | --- | --- | --- |
| Intercept | -0.083218 | 0.076251 | -1.0914 | 0.27538 | -0.23285 | 0.066412 |  |
| oneBackSameGem | 0.62289 | 0.19375 | 3.2149 | 0.0013463 | 0.24269 | 1.0031 | *** |
| twoBackSameGem | 0.29632 | 0.12814 | 2.3126 | 0.020948 | 0.044877 | 0.54777 | * |
| threeBackSameGem | 0.35279 | 0.13314 | 2.6498 | 0.008181 | 0.091526 | 0.61405 | ** |
| fourBackSameGem | 0.32482 | 0.13215 | 2.4579 | 0.014142 | 0.065493 | 0.58414 | * |
| fiveBackSameGem | 0.23133 | 0.12847 | 1.8006 | 0.07206 | -0.020773 | 0.48343 |  |
| oneBackOtherGem | 0.074356 | 0.10218 | 0.72769 | 0.46697 | -0.12616 | 0.27487 |  |
| twoBackOtherGem | 0.08851 | 0.1104 | 0.8017 | 0.42292 | -0.12814 | 0.30516 |  |
| threeBackOtherGem | -0.17593 | 0.10473 | -1.6798 | 0.093307 | -0.38144 | 0.029589 |  |
| fourBackOtherGem | -0.17049 | 0.11637 | -1.465 | 0.14323 | -0.39885 | 0.057872 |  |
| fiveBackOtherGem | 0.12256 | 0.11191 | 1.0951 | 0.27373 | -0.097052 | 0.34217 |  |

**Supplementary Table 3.** Mean (standard deviation) number of trials in each condition for the RSA conducted at time of door onset presented in the main paper. There is no significant difference between the mean trial numbers per bin in dependent and independent condition (t(28 = 1.50, p = 0.55, 95% CI [-.02, .15]). This was calculated by comparing the mean trial numbers in each probability bin of the two contexts in the dependent (8 probability bins total) to those in the independent condition using a paired samples ttest.

|  | Dependent Condition |  | Independent Condition |  |
| --- | --- | --- | --- | --- |
|  | Context 1 | Context 2 | Context 3 | Context 4 |
| Quartile 1 | 16.24<br>(3.07) | 16.17<br>(2.51) | 15.55<br>(4.20) | 15.14<br>(3.03) |
| Quartile 2 | 15.97<br>(3.12) | 15.45<br>(1.86) | 15.14<br>(3.66) | 16.45<br>(3.17) |
| Quartile 3 | 15.07<br>(2.40) | 15.55<br>(3.44) | 15.93<br>(4.25) | 16.38<br>(3.70) |
| Quartile 4 | 16.00<br>(3.26) | 15.69<br>(2.95) | 16.21<br>(3.92) | 14.83<br>(3.67) |

**Supplementary Table 4.** Statistics for the crossentropy loss bins at time of onset (crossvalidation, one-hot)

| Bins | 1 | 2 | 3 | 4 | 5 | 6 | 7 | 8 | 9 | 10 |
| --- | --- | --- | --- | --- | --- | --- | --- | --- | --- | --- |
| T-value | -0.45 | -0.03 | -1.53 | -2.12 | -2.39 | -3.12 | -2.41 | -1.72 | -2.53 | -2.60 |
| P value | 0.33 | 0.49 | 0.07 | 0.02 | 0.01 | 0.002 | 0.01 | 0.05 | 0.009 | 0.007 |

**Supplementary Figure 1.**
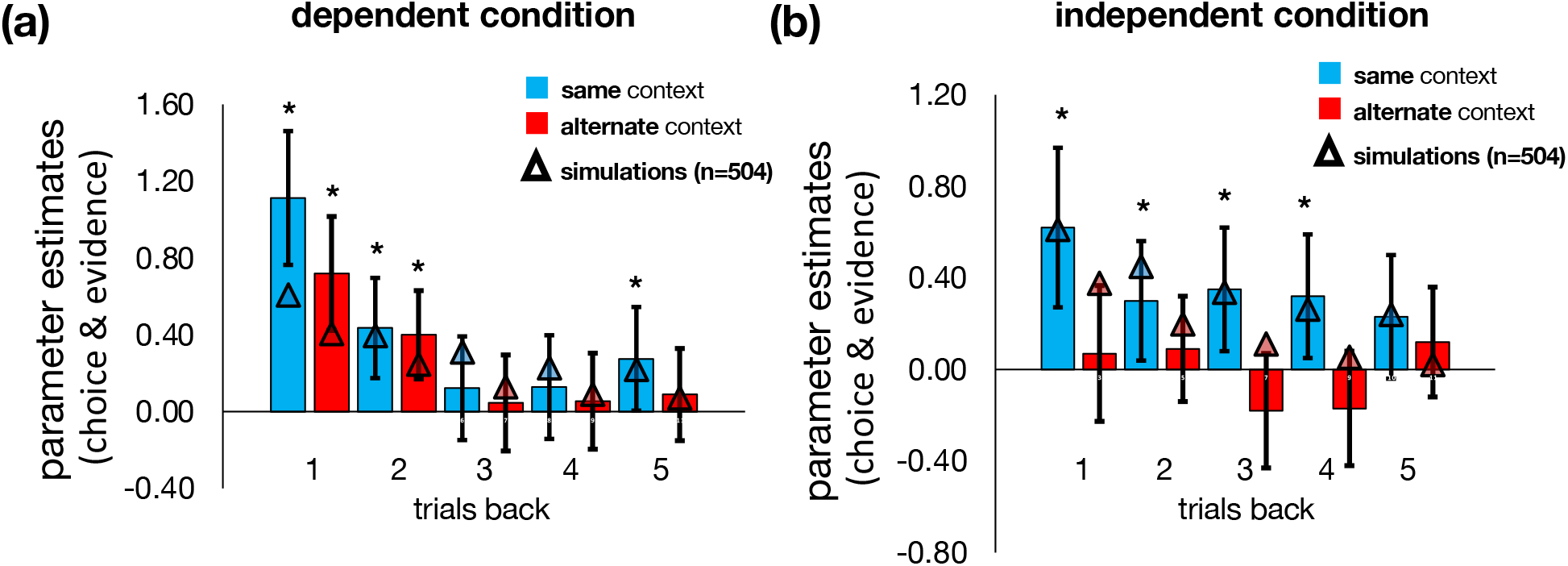
Behavioural data. Parameter estimates predicting choice from state transitions 1-5 trials back (same data as in Figure 2) with simulations from the fixed model (triangles) superimposed. Bars represent fixed effect regression coefficients from a mixed effects logistic regression. Error bars represent standard error of the mean. *p<0.05.

**Supplementary Figure 2.**
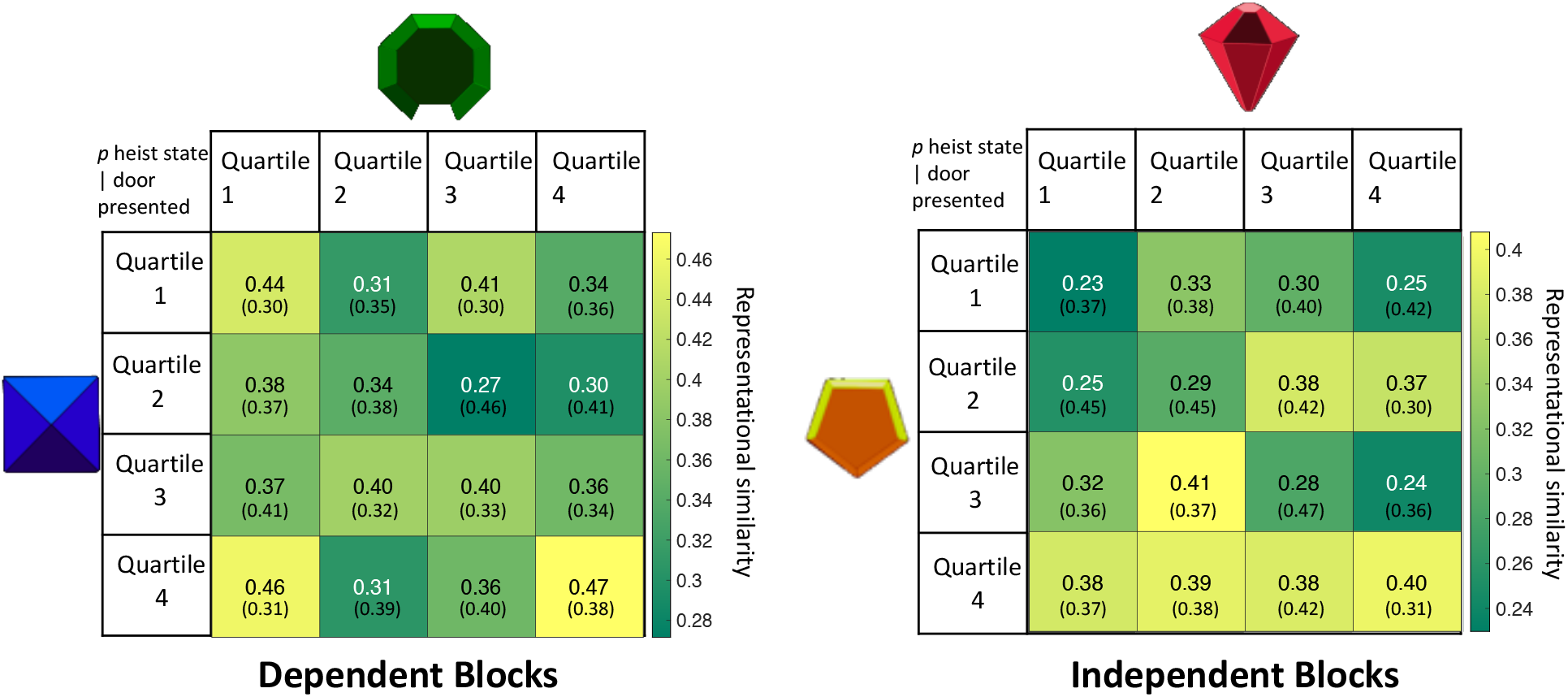
RSA at door onset. Each entry shows the mean (standard deviation) similarity score over participants 4*4 quartiles of p (heist state | door presented). Analysis in the main text averages on and off diagonal cells to generate a matched and mismatched similarity score for each condition (see Figure 3c main text). There was a significant interaction between condition and on and off-diagonal (t(28) = 4.02, p < 0.001, 95% CI [.11,.33]), characterised by a significant difference in similarity between on and off diagonal (Figure 3c) in the dependent condition (t(28) = 5.33, p < 0.001, 95% CI [.11,.26]) which was not observed in the independent condition (t(28) = −0.82, p = 0.42, 95% CI [-.11, .05]). As reported in-text, there was no difference between trial numbers in the two conditions (t(28 = 1.50, p = 0.55, 95% CI [-.02, .15]), calculated by comparing the mean number of trials in dependent and independent blocks’ probability bins (two-tailed t-test).

**Supplementary Table 5.** Paired comparisons of the interaction effect at onset timepoint across the anatomical subsections of the MTL shown in Figure 4a. As we have 6 paired comparisons, the Bonferroni-corrected threshold for significance is 0.05/6 = 0.0083.

| Interaction values for regions: | Mean difference (SD) | SEM | t-value | pValue (2-tailed) | (95% CIs): |  |
| --- | --- | --- | --- | --- | --- | --- |
| Hippocampus – entorhinal cortex | 0.10 (0.25) | 0.05 | 2.07 | 0.048 | 0.00 | 0.19 |
| Hippocampus - amygdala | 0.10 (0.23) | 0.04 | 2.27 | 0.031 | 0.01 | 0.19 |
| Hippocampus - parahippocampus | 0.13 (0.18) | 0.03 | 3.91 | 0.001 | 0.06 | 0.19 |
| Entorhinal cortex - amygdala | -0.00 (0.21) | 0.04 | -0.00 | 0.988 | -0.08 | 0.08 |
| Entorhinal cortex - parahippocampus | 0.03 (0.17) | 0.03 | 0.92 | 0.366 | -0.04 | 0.09 |
| Amygdala - parahippocampus | 0.03 (0.23) | 0.04 | 0.70 | 0.490 | -0.06 | 0.12 |

**Supplementary Figure 3.**
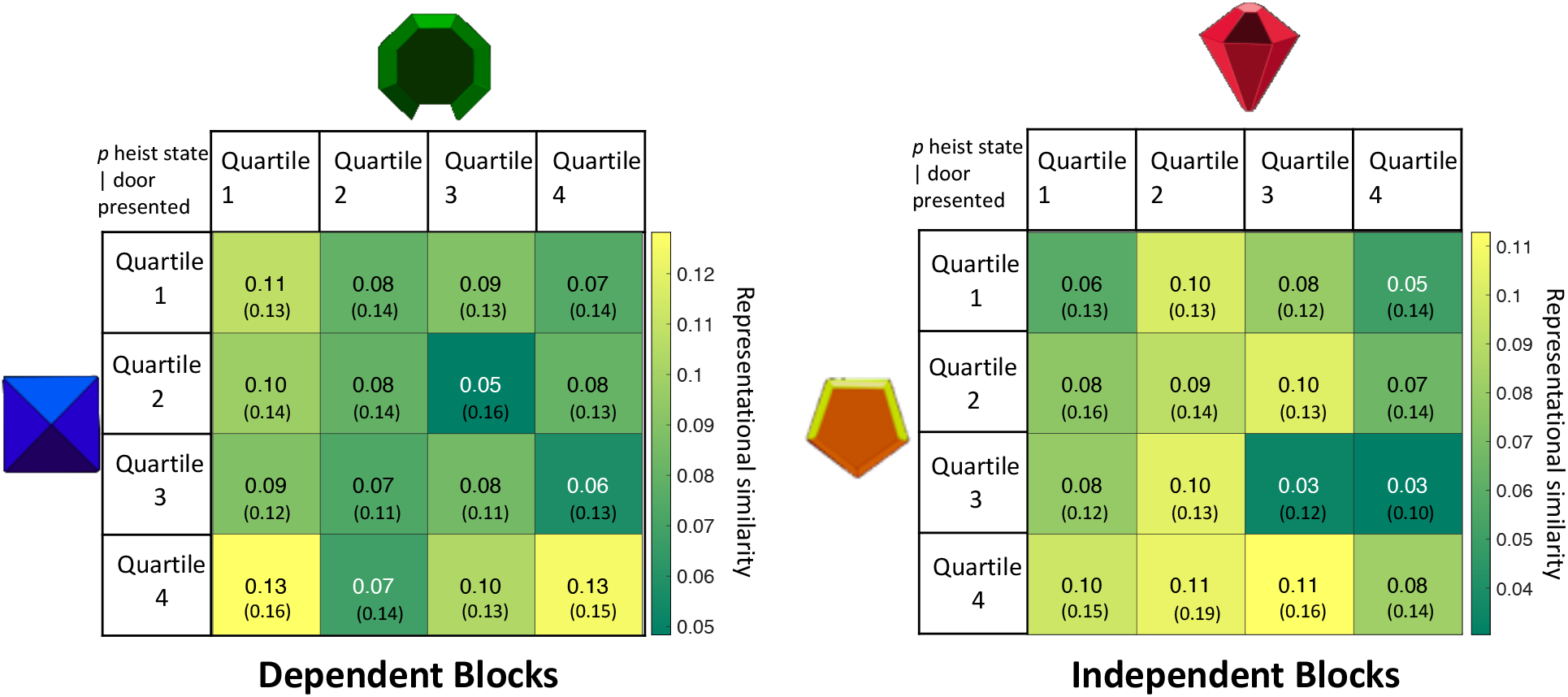
RSAs with 4*4 quartiles in cross validation. Here, the similarity between bin *n*_*i*_ and *n*_*j*_ where *i* and *j* are drawn from different scanner runs. Condition × bin interaction: t(28) = 1.89, p = 0.068; 95% CI [-.00, .07]; Matched vs mismatched in dependent condition: t(28) = 1.79; p = 0.084; 95% CI [-.00,.04]; Matched vs mismatched in independent condition: t(28) = −1.35; p = 0.188; 95% CI [-.04, .01].

**Supplementary Figure 4.**
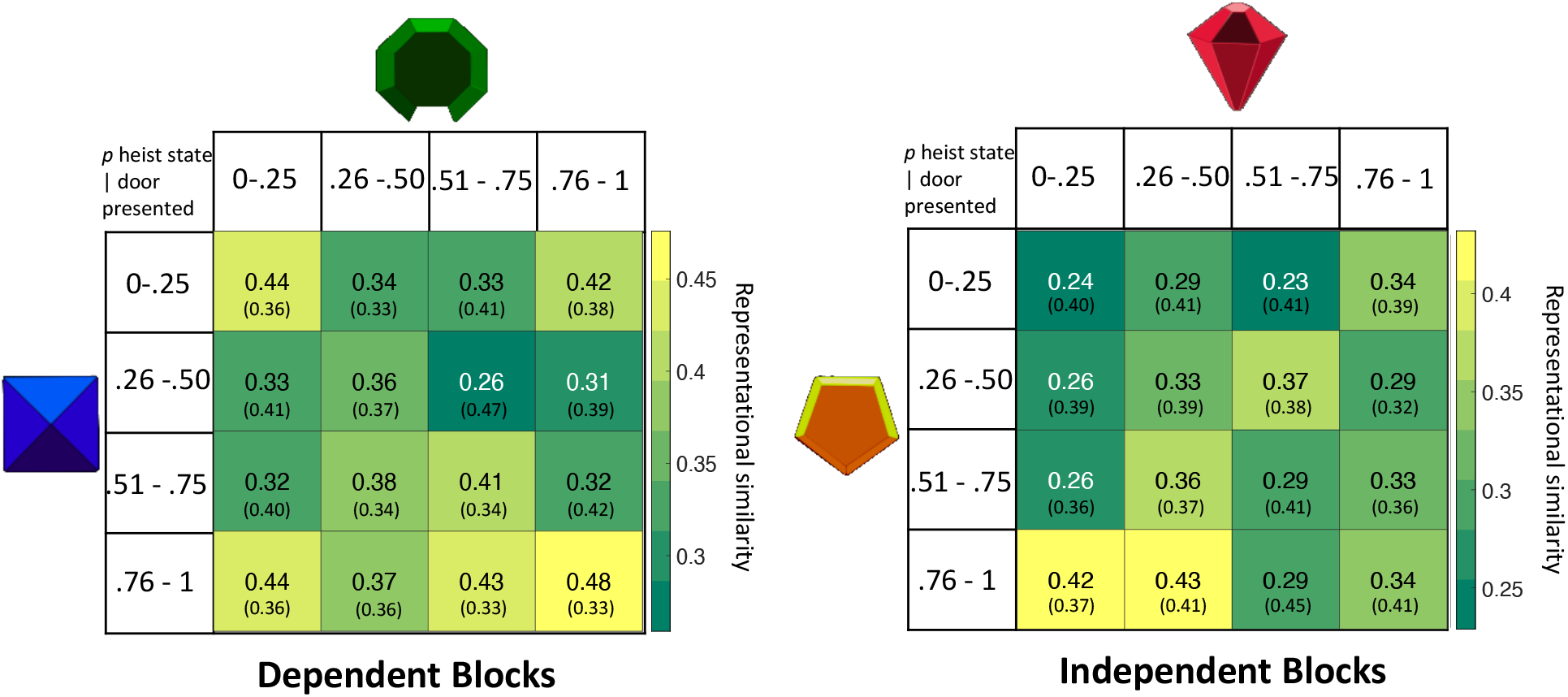
Same RSA analysis as reported in the main text and Supplementary Figure 2, now using fixed range of quartiles (0-0.25, 0.26-0.5, 0.51-0.75 and 0.76-1.0) to compute similarity scores. There was a significant condition by diagonal interaction (dependent, independent) × bin (on, off diagonal) (t(28) = 4.20, p < 0.001; 95% CI [.10, .30]), characterised by a significant difference in similarity between on and off diagonal (Figure 3c) in the dependent condition (t(28) = 5.73; p < 0.001; 95% CI [.13, .28]) which was not observed in the independent condition (t(28) = 0.04; p = 0.97; 95% CI [-.07, .08]).

**Supplementary Figure 5.**
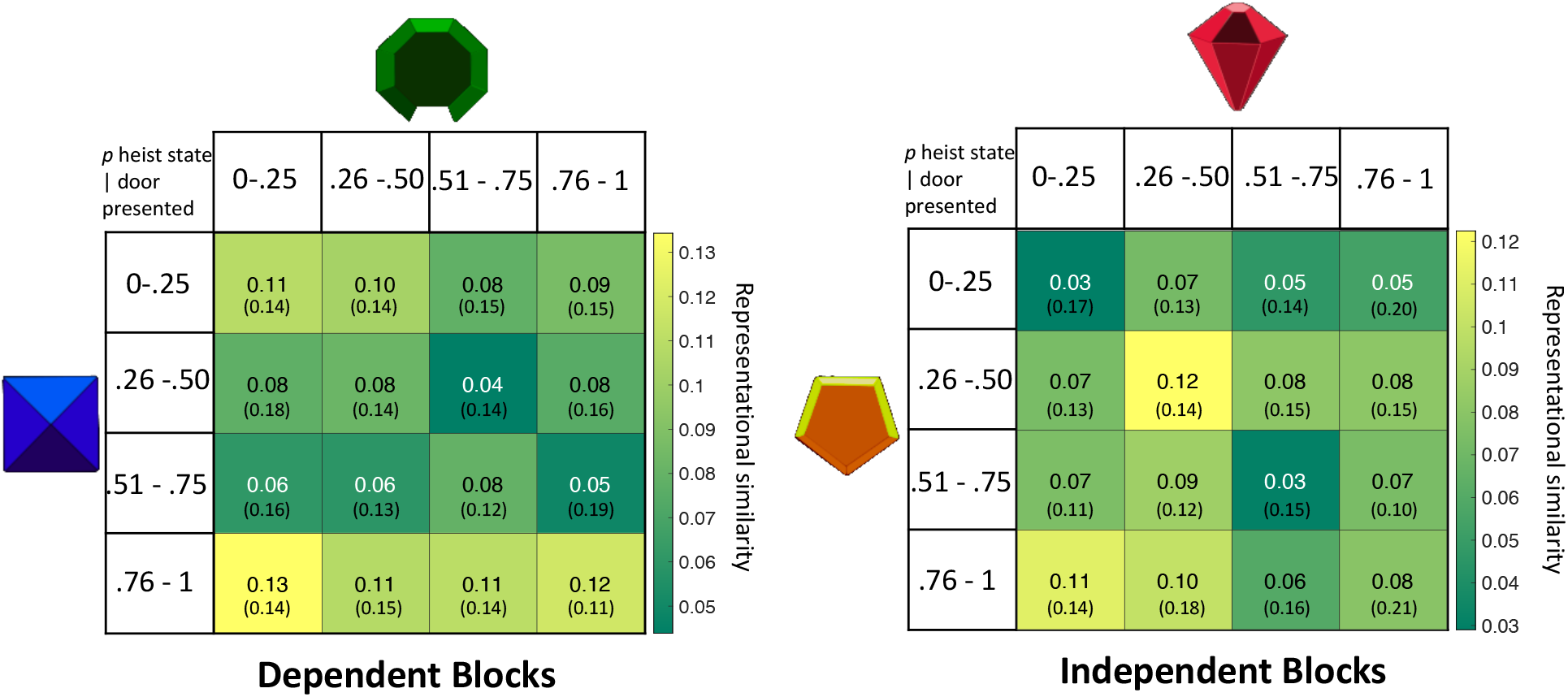
Same RSA analysis as in Supplementary Figure 4 in crossvalidation. Condition by diagonal interaction: t(28) = 2.04, p = 0.05; 95% CI [.00, .06]. Similarity between on and off diagonal (dependent condition: t(28) = 1.22; p = 0.23; 95% CI [-.01, .04], independent condition: t(28) = −1.34; p = 0.19; 95% CI [-.04, .01]).

**Supplementary Figure 6.**
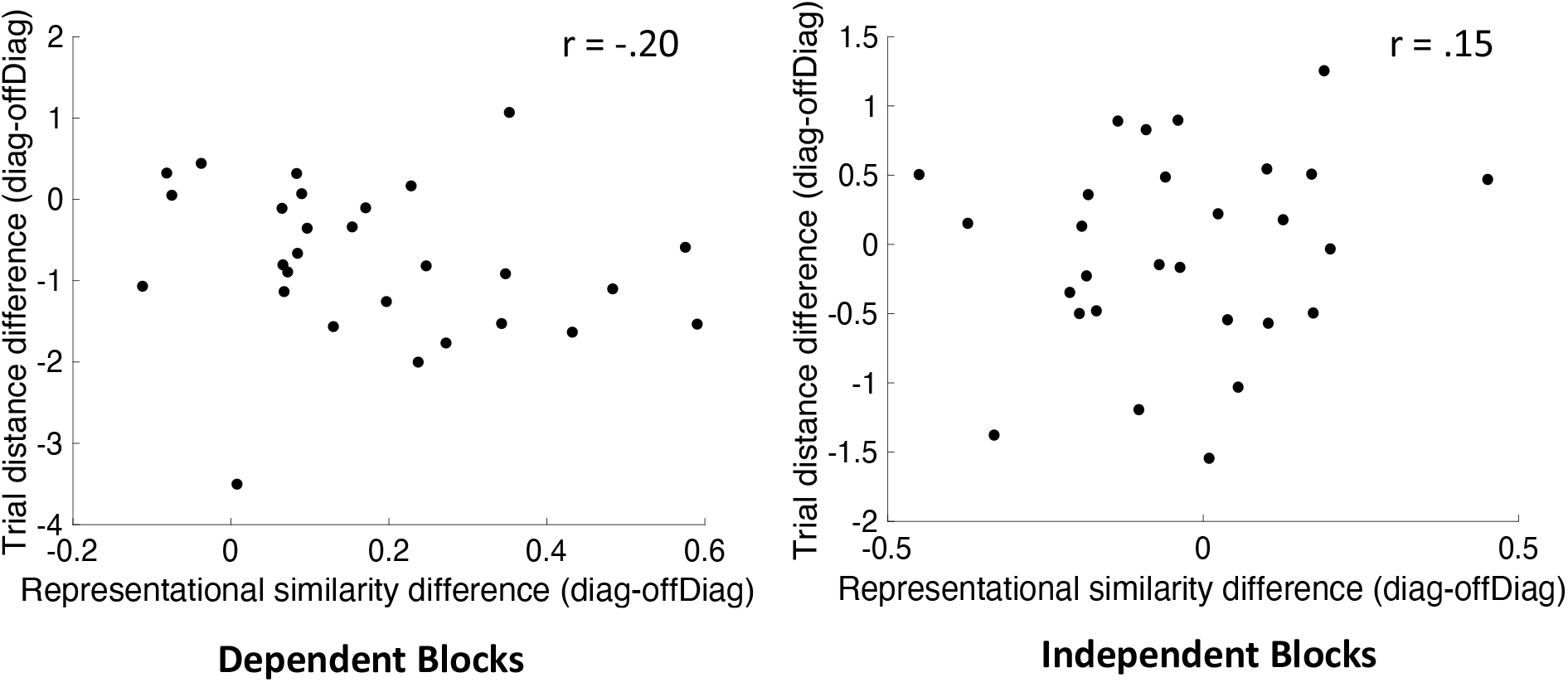
Scatterplots showing the relationship between the difference in mean trial distance (temporal distance) between the diagonal and off-diagonal trials and the representational similarity difference between diagonal and off-diagonal (see **Figure 3** and **Supplementary Figure 2**) at the time of door onset for the two conditions.

**Supplementary Figure 7.**
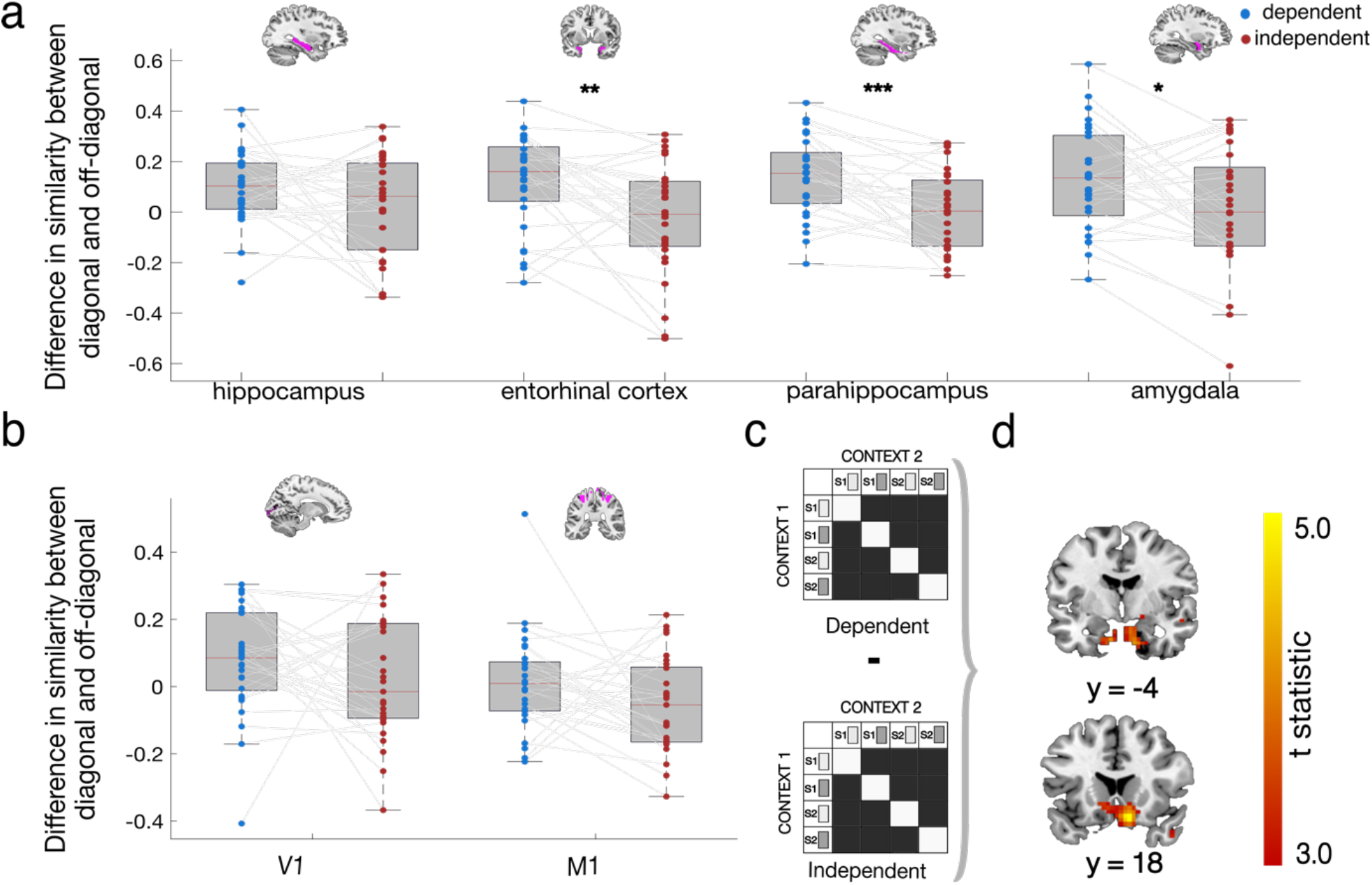
**(a)** To investigate whether the effect observed at the time of feedback (Main Text Figure 5) was specific to a particular subregion of the MTL we conducted follow-up RSAs in the same subregions of the MTL as the time of door onset. Difference in similarity scores (diagonal minus off diaganal) were entered into a 2(condition) by 4(region) repeated measures ANOVA. This found no evidence for functional specialisation within the MTL (no condition*region interaction: F(3,81) = 1.21, p = 0.312) and no main effect of region (no main effect of region F(3,81) = 0.95, p = 0.419). However there was a main effect of condition. In other words, across all regions, the similarity of identical compared to non-identical action-outcome state pairings was greater in the dependent than the independent condition (F(1,27) = 10.06, p = 0.004). **(b)** We then repeated this analysis in two control regions (V1 and M1) and found no evidence for differential encoding of probability bins between conditions in either region (V1: interaction t(28) = 1.16, p = 0.26, 95% CI [-.04,.16]; M1: interaction t(28) = 1.66, p = 0.11, 95% CI [-.02,.15]). **C)** A whole brain searchlight comparing the difference in similarity scores between on and off diagonal between conditions revealed a significant interaction within our state prediction error ROI (left: [-16, −4, −32], t(28) = 3.47, p = 0.02 cluster level corrected within our SPE mask), as well as in a cluster comprised of right hippocampus extending into pons (t(28) = 5.05, p < 0.0001 uncorrected, [12, 18, −18], p = 0.04 cluster level corrected across the whole brain with cluster-forming threshold of *p* < .001 uncorrected). No other significant effects were observed.

**Supplementary Figure 8.**
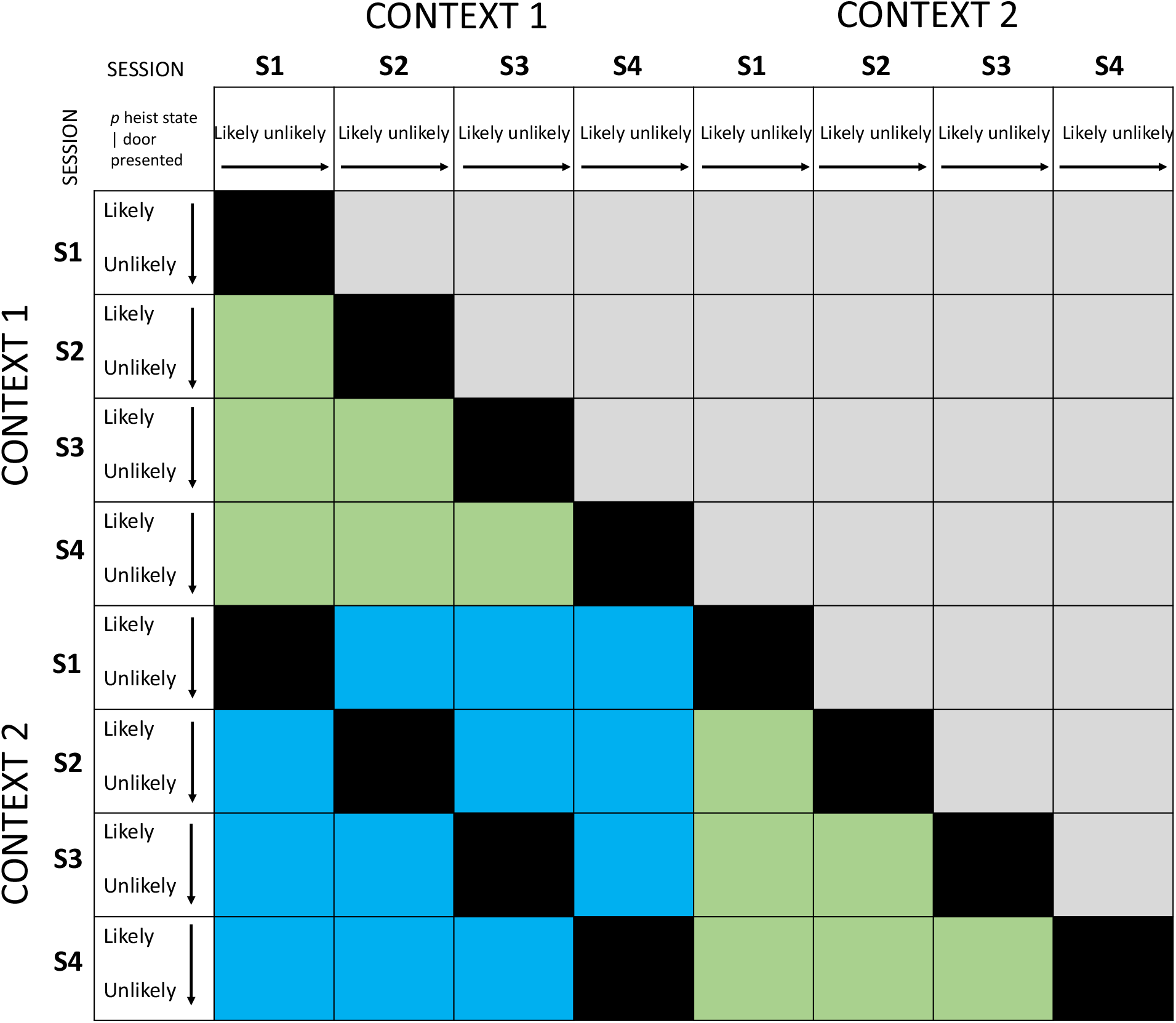
Illustration of how the crossvalidation was implemented. Each square corresponds to a 4×4 RSM of probability bins’ representational similarity to each other. To determine the between-context difference in similarity between the diagonal and off-diagonal in crossvalidation, we averaged across all Fisher transformed between-context similarity squares that are indicated in blue, i.e. that were not part of the same scanner session (e.g. c_1_ S1 to c_2_ S2) and then computed the statistics on the resulting averaged 4×4 as described in-text for the main analysis. To compute the same – same context crossvalidated similarity shown in supplementary Figure 5, we averaged across all Fisher transformed similarity squares indicated in green and computed the statistics as described for the main analysis in-text.

**Supplementary Figure 9.**
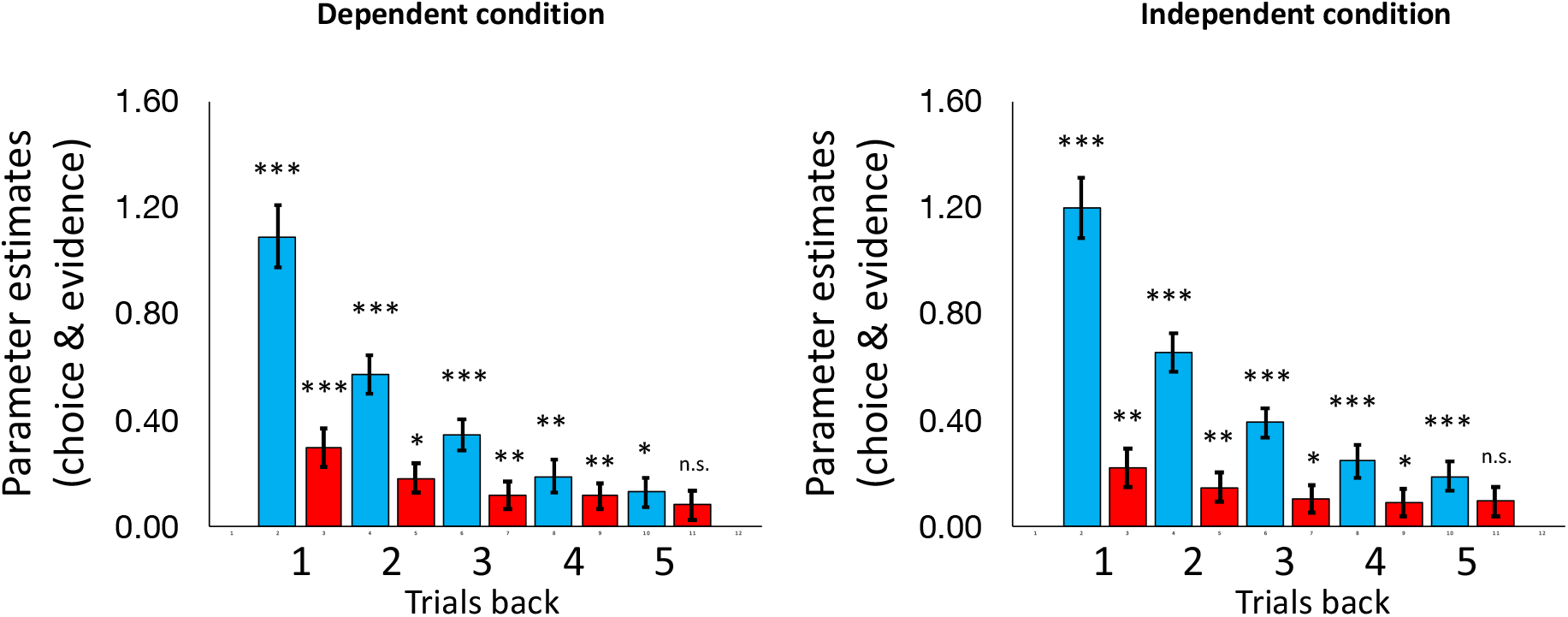
Parameter estimates predicting choice from state transitions 1-5 trials back from a separate behavioural experiment in which conditions were not signalled to participants (same analysis as shown Figure 2 Main Paper). Bars represent fixed effect regression coefficients from a mixed effects logistic regression (see also **Supplementary Table 6** and **Table 7** below). Error bars represent standard error of the mean. *p < 0.05, ** p < 0.01, *** p < 0.001. Participants integrate evidence from the other context in the dependent condition and in the independent condition. As in **Supplementary Figure 1**, blue signals information from the same context and red from the other context.

**Supplementary Table 6.** Dependent Condition (separate behavioural experiment in which conditions were not signalled to participants during the experiment and participants were not told about them in advance): Fixed effect coefficients from Lagged Logistic Regression Model (N=31).*p<0.05, **p<0.01, ***, p<0.001

| (95% CIs): |  |  |  |  |  |  |  |
| --- | --- | --- | --- | --- | --- | --- | --- |
| Name | Estimate | SE | tstat | pValue | Lower | Upper |  |
| Intercept | -0.030993 | 0.031728 | -0.97682 | 0.3286917 | -0.093189 | 0.03120381 |  |
| oneBackSameGem | 1.0902467 | 0.114242 | 9.543293 | 1.85E-21 | 0.866299 | 1.31419453 | *** |
| twoBackSameGem | 0.570577 | 0.072219 | 7.900604 | 3.18E-15 | 0.429006 | 0.71214793 | *** |
| threeBackSameGem | 0.3455804 | 0.056242 | 6.144565 | 8.44E-10 | 0.2353305 | 0.45583036 | *** |
| fourBackSameGem | 0.1868641 | 0.061791 | 3.024112 | 0.0025023 | 0.0657351 | 0.30799314 | ** |
| fiveBackSameGem | 0.1284043 | 0.056651 | 2.266604 | 0.0234436 | 0.0173529 | 0.23945568 | * |
| oneBackOtherGem | 0.2962969 | 0.072803 | 4.069852 | 4.75E-05 | 0.1535822 | 0.43901153 | *** |
| twoBackOtherGem | 0.1817307 | 0.055763 | 3.258962 | 0.0011233 | 0.0724183 | 0.291043 | * |
| threeBackOtherGem | 0.1148956 | 0.051048 | 2.250754 | 0.0244308 | 0.0148275 | 0.21496379 | ** |
| fourBackOtherGem | 0.114004 | 0.050795 | 2.244416 | 0.0248356 | 0.014432 | 0.21357593 | ** |
| fiveBackOtherGem | 0.0808322 | 0.054348 | 1.48731 | 0.1369761 | -0.025705 | 0.18736991 |  |

**Supplementary Table 7.**
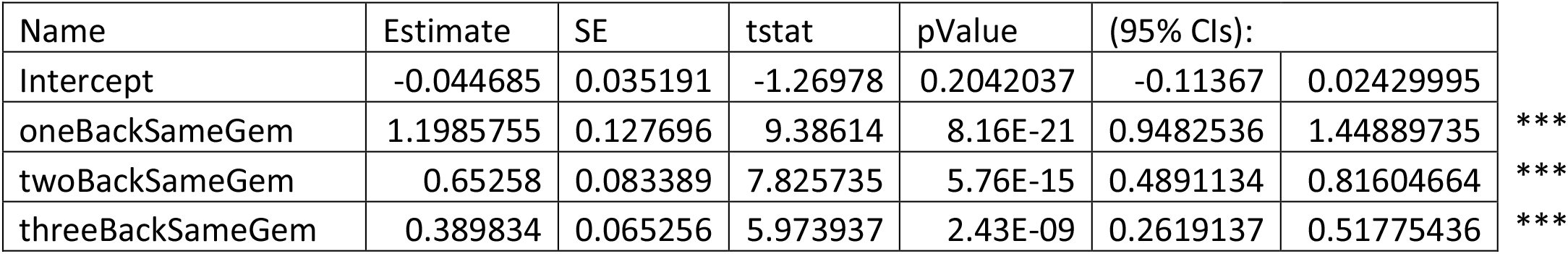
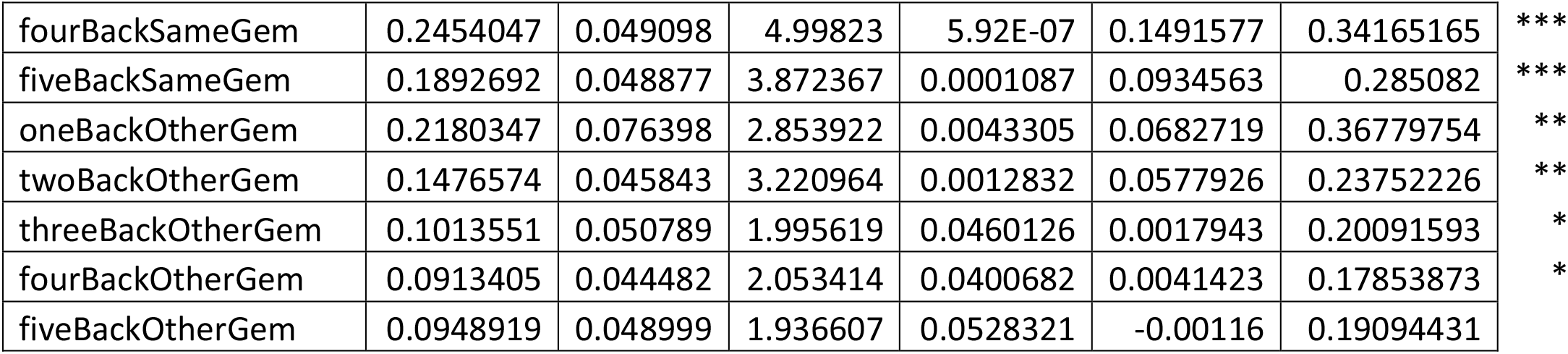
Independent Condition (separate behavioural experiment in which conditions were not signalled to participants during the experiment and participants were not told about them in advance): Fixed effect coefficients from Lagged Logistic Regression Model (N=31).*p<0.05, **p<0.01, ***, p<0.001

| Name | Estimate | SE | tstat | pValue | (95% CIs): |  |
| --- | --- | --- | --- | --- | --- | --- |
| Intercept | -0.044685 | 0.035191 | -1.26978 | 0.2042037 | -0.11367 | 0.02429995 |
| oneBackSameGem | 1.1985755 | 0.127696 | 9.38614 | 8.16E-21 | 0.9482536 | 1.44889735 |
| twoBackSameGem | 0.65258 | 0.083389 | 7.825735 | 5.76E-15 | 0.4891134 | 0.81604664 |
| threeBackSameGem | 0.389834 | 0.065256 | 5.973937 | 2.43E-09 | 0.2619137 | 0.51775436 |

|  |  |  |  |  |  |  |  |
| --- | --- | --- | --- | --- | --- | --- | --- |
| fourBackSameGem | 0.2454047 | 0.049098 | 4.99823 | 5.92E-07 | 0.1491577 | 0.34165165 | *** |
| fiveBackSameGem | 0.1892692 | 0.048877 | 3.872367 | 0.0001087 | 0.0934563 | 0.285082 | *** |
| oneBackOtherGem | 0.2180347 | 0.076398 | 2.853922 | 0.0043305 | 0.0682719 | 0.36779754 | ** |
| twoBackOtherGem | 0.1476574 | 0.045843 | 3.220964 | 0.0012832 | 0.0577926 | 0.23752226 | ** |
| threeBackOtherGem | 0.1013551 | 0.050789 | 1.995619 | 0.0460126 | 0.0017943 | 0.20091593 | * |
| fourBackOtherGem | 0.0913405 | 0.044482 | 2.053414 | 0.0400682 | 0.0041423 | 0.17853873 | * |
| fiveBackOtherGem | 0.0948919 | 0.048999 | 1.936607 | 0.0528321 | -0.00116 | 0.19094431 |  |

## Notes

### Competing Interest Statement

The authors have declared no competing interest.

### Summary of Updates

Funding information was incomplete in previous version

